# Clathrin-mediated Endocytosis Facilitates Internalization of *Magnaporthe oryzae* Effectors into Rice Cells

**DOI:** 10.1101/2021.12.28.474284

**Authors:** Ely Oliveira-Garcia, Tej Man Tamang, Jungeun Park, Melinda Dalby, Magdalena Martin-Urdiroz, Clara Rodriguez Herrero, An Hong Vu, Sunghun Park, Nicholas J. Talbot, Barbara Valent

## Abstract

Fungi and oomycetes deliver effectors into living plant cells to suppress defenses and control plant processes needed for infection. Little is known about the mechanism by which these pathogens translocate effector proteins across the plasma membrane into the plant cytoplasm. The blast fungus *Magnaporthe oryzae* secretes cytoplasmic effectors into a specialized biotrophic interfacial complex (BIC) before translocation. Here we show that cytoplasmic effectors within BICs are packaged into vesicles that are occasionally observed in the host cytoplasm. Live cell imaging with fluorescently-labeled rice showed that effector vesicles colocalize with plant plasma membrane and with clathrin light chain-1, a marker for clathrin-mediated endocytosis (CME). Inhibition of CME using Virus-Induced Gene Silencing (VIGS) and chemical treatments results in cytoplasmic effectors in swollen BICs lacking vesicles. In contrast, fluorescent marker co-localization, VIGS and chemical inhibitor studies failed to support a major role for clathrin-independent endocytosis in effector vesicle formation. Localization studies of two novel effectors, and of known effectors after CME inhibition, indicate that cytoplasmic effector translocation occurs underneath appressoria before invasive hyphal growth. Taken together, this study provides evidence that cytoplasmic effector translocation is mediated by clathrin-mediated endocytosis in BICs and suggests a role for *M. oryzae* effectors in co-opting plant endocytosis.

## INTRODUCTION

Many filamentous eukaryotic plant pathogens, such as fungi and oomycetes, cause plant disease by hijacking and feeding on living plant cells, and they deliver effectors into and around host cells to promote infection (Giraldo and Valent, 2013; Lo Presti and Kahmann, 2017). This includes the Ascomycete fungus *Magnaporthe oryzae* (synonym of *Pyricularia oryzae*), which threatens global food security by causing blast diseases on rice, on several millets, and most recently on wheat (Gladieux et al., 2018; Valent et al., 2020). *M. oryzae* executes a hemibiotrophic lifestyle involving biotrophic invasion of successive plant cells by specialized intracellular invasive hyphae (IH) that grow from cell to cell (Kankanala et al., 2007). Blast IH growing in living host cells are enclosed by extensions of the plant plasma membrane termed the extra-invasive hyphal membrane (EIHM). Colonized cells die as they fill with hyphae and the EIHM and host vacuolar membranes become disrupted. Meanwhile, the fungus moves on to colonize neighboring cells using the same invasion strategy (Kankanala et al., 2007; Mochizuki et al., 2015; Sakulkoo et al., 2018; Jones et al., 2021).

Live cell imaging of blast IH invading optically clear rice leaf sheath cells has defined a dimorphic hyphal switch associated with secretion and targeting of effectors – small, secreted proteins that suppress immunity responses in host plant cells. Specifically, cytoplasmic effectors, destined for translocation inside the host cell, are associated with the first two IH cells to grow in the plant cell lumen, the tubular primary invasive hypha that grows directly after penetration and the pseudohyphal bulbous IH cells that subsequently grow to fill the host cell (Kankanala et al., 2007; Khang et al., 2010). Effectors destined for delivery to the host cytoplasm are secreted by a specialized, Brefeldin A (BFA)-insensitive, Golgi-independent secretion system (Giraldo et al., 2013). After secretion, these ‘cytoplasmic effectors’ accumulate in an extended dome-shaped interfacial region of the EIHM matrix, the Biotrophic Interfacial Complex (BIC). BICs first occur as ‘tip-BICs’ at primary hyphal tips, and then as ‘side-BICs’ that move beside the first bulbous IH cell (Khang et al., 2010). During live cell imaging, fluorescent cytoplasmic effectors are routinely detected in the cytoplasm and/or nuclei of invaded plant cells. Some cytoplasmic effectors are also detected in the cytoplasm of surrounding plant cells, apparently moving through plasmodesmata to prepare neighboring cells before invasion (Khang et al., 2010). In contrast, apoplastic effectors, such as the biotrophy-associated secreted protein 4 (Bas4), the LysM protein 1 (Slp1), and Bas113, are secreted via conventional Golgi-dependent secretion, and accumulate in the EIHM around the entire IH (Giraldo et al., 2013). The retention of apoplastic effectors such as Bas4 inside the EIHM (Mosquera et al., 2009; Khang et al., 2010), lack of separation of the EIHM from IH during plasmolysis (Kankanala et al., 2007), and exclusion of the endocytosis tracker dye FM4-64 from IH membranes inside the FM4-64 stained EIHM (Kankanala et al., 2007; Giraldo et al., 2013), all indicate that IH grow inside a sealed EIHM compartment.

The mechanisms by which eukaryotic filamentous plant pathogens deliver effectors across the host plasma membrane into living host cells are poorly understood, and may be conserved or evolutionarily unique for different host-pathogen interactions (Giraldo and Valent, 2013; Petre and Kamoun, 2014; Fawke et al., 2015; Lo Presti et al., 2015; Lo Presti and Kahmann, 2017). The oomycete potato pathogen *Phytophthora infestans* resembles *M. oryzae* in having distinct secretion systems for different classes of effectors. That is, cytoplasmic effectors are secreted by a Brefeldin A (BFA)-insensitive (Golgi-independent) secretion and apoplastic effectors are secreted by conventional BFA-sensitive secretion (Giraldo et al., 2013; Van den Ackerveken, 2017; Wang et al., 2017). Although cell entry motifs for fungal effectors have not been identified, oomycete effectors contain amino acid translocation motifs, with the RXLR motif being the most common. The RXLR motif in oomycete effectors was reported to mediate host cell uptake through endocytosis based on binding to phosphatidylinositol-3-phosphate (PI3P) in host membrane (Kale et al., 2010), although it was subsequently shown that the RXLR motif from *P. infestans* effector AVR3a is cleaved off before secretion from the pathogen (Wawra et al., 2017). Therefore, the function of RXLR motif in cell entry remains disputed (Petre and Kamoun, 2014; Trusch et al., 2018). Another amino acid motif, YKARK, appears involved in translocation of the effector SpHtp3 from *Saprolegnia parasitica*, a serious oomycete pathogen of fish, inside fish cells (Trusch et al., 2018). Specifically, this effector appears to be internalized inside host cells via lipid-raft mediated endocytosis through binding to a gp96-like receptor, and another effector is involved in releasing SpHtp3 from vesicles into the fish cytoplasm. In an example from a fungal pathogen, a solvent-exposed RGD vitronectin-like motif is required for internalization of ToxA, a host-specific proteinaceous toxin secreted by the necrotrophic wheat pathogen *Pyrenophora tritici-repentis* (Manning and Ciuffetti, 2005). In addition to uptake by plant endocytosis, effectors might enter plant cells through specialized translocon complexes in the plasma membrane. For example, the human malaria parasite, *Plasmodium falciparum*, is contained within a parasitophorous vacuole inside invaded red blood cells, and effectors are delivered across the membrane into the host cytoplasm by the PTEX translocon (Elsworth et al., 2014). Recently, a stable protein complex comprised of five unrelated fungal effectors and two fungal membrane proteins has been implicated in effector translocation by the maize smut fungus *Ustilago maydis* (Ludwig et al., 2021). However, the general question of how effectors are taken up into plant cells remains largely unresolved for eukaryotic plant pathogens.

Clathrin-mediated endocytosis (CME) is the major mechanism by which eukaryotic cells internalize extracellular or membrane-bound cargoes, and this process is best understood in mammalian systems and yeast (Kaksonen and Roux, 2018). Increasing understanding of CME in plants shows both conserved and evolutionarily unique mechanistic details (Qi et al., 2018; Reynolds et al., 2018; Dejonghe et al., 2019; Johnson et al., 2020; Narasimhan et al., 2020). CME is known to play key roles in plant-microbe interactions (Dodds and Rathjen, 2010; Beck et al., 2012; Fan et al., 2015). CME is required for immunity mediated by pattern recognition receptor kinases, for instance, specifically through internalization of activated extracellular pattern recognition receptors (PRRs) for degradation in the vacuole (Mbengue et al., 2016). Examples include the pattern recognition receptor Pep receptor 1/2 (PEPR1/2), the EF-TU receptor (EFR) recognizing the PAMP bacterial elongation factor, and FLS2 recognizing the PAMP flagellin (flg-22) (Spallek et al., 2013; Mbengue et al., 2016), as well as the Cf-4 receptor-like protein that recognizes the *C. fulvum* Avr4 avirulence effector (Postma et al., 2016). Trafficking of activated PRRs requires clathrin and converges into endosomal vesicles that are also shared by the hormone receptor Brassinosteriod Insensitive 1 (BRI1). Additionally, it is known that pathogens can control plant CME. In addition to its role in suppressing INF1-mediated cell death, the *P. infestans* host-translocated RXLR effector AVR3a was recently shown to associate with the Dynamin-Related Protein 2, a plant GTPase implicated in receptor-mediated endocytosis, and to suppress flg22-triggered defense responses and reduce FLS2 internalization (Chaparro-Garcia et al., 2015). Therefore, AVR3a interacts with a membrane-remodeling complex involved in immune receptor-mediated endocytosis. Also, in host cells containing haustoria from the oomycete *P. infestans*, the late endosomal pathway is rerouted from the vacuole to the extrahaustorial membrane (EHM) surrounding haustoria (Bozkurt et al., 2015). Multiple effectors from this pathogen are now implicated in rerouting the host membrane trafficking system and autophagy machinery for the benefit of the invading pathogen (Bozkurt et al., 2015; Dagdas et al., 2018; Petre et al., 2021).

Effector translocation in the *M. oryzae*/rice pathosystem can be assayed directly by live cell imaging of fluorescently-labeled effector proteins inside host cells after secretion by IH invading rice leaf sheath cells (Khang et al., 2010). Although some effectors naturally localize to host nuclei, addition of a nuclear localization signal (NLS) to fluorescent effectors enhances visualization of translocation and cell-to-cell movement. Blast cytoplasmic effectors apparently do not contain a translocation motif as reported for oomycete effectors. Instead, preferential accumulation in the BIC is the major predictor for translocation into the rice cell. Previously, Nishizawa and colleagues reported that a cytoplasmic effector occurs in punctae (vesicles) in an outer BIC region (Mochizuki et al., 2015; Nishimura et al., 2016; Nishizawa et al., 2016). Here, we have used live-cell fluorescence imaging to characterize the nature and dynamics of membrane-bound effector vesicles within BICs and in surrounding host cytoplasm. We show that BICs are enriched for CME components, and we report functional analyses using virus-induced gene silencing (VIGS) and pharmacological approaches. When considered together, our data indicate that blast effectors are internalized through host clathrin-mediated endocytosis in the outer BIC layers, followed by escape from the vesicles into the rice cytoplasm.

## RESULTS

### BICs contain multiple vesicles labeled by cytoplasmic effectors

BICs were previously reported as interfacial structures containing fluorescently-labelled apoplastic effectors, such as Bas4:eGFP, in an inner base layer and a concentration of fluorescent cytoplasmic effectors in the outer ‘dome-shaped’ region (Khang et al., 2010; Nishizawa et al., 2016). Using laser confocal imaging, we now routinely resolve the dome region of BICs as a concentration of vesicles containing fluorescent cytoplasmic effectors (Figure 1). These effector vesicles are illustrated by an image of a BIC at the tip of a primary hypha (tip-BIC) expressing fluorescent apoplastic effector Bas4:eGFP and fluorescent cytoplasmic effector Pwl2:mRFP (Figure 1A). The Bas4:eGFP fluorescence remains distributed throughout the EIHM matrix surrounding the entire growing IH, including in an inner layer of the BIC. In contrast, Pwl2:mRFP occurs as distinct vesicles in the outer BIC region. We have observed effector vesicles in tip-BICs labeled with additional translocated effectors Bas1 and Pwl1 fused to either eGFP or mRFP (Supplemental Figure S1). Vesicles are also observed after tip-BICs move beside the first bulbous IH cells (side-BICs) (Figure 1B; Supplemental Figure S2). We evaluated the proximity of these vesicles relative to both tip- and side-BICs. Whereas the vast majority of vesicles associated with primary IH were located in the tip-BIC, some vesicles associated with side-BICs were found in regions of the cytoplasm at least 7 µm from the fungal cell wall at the BIC, which we define as outside the BIC (Figure 1C). Inclusion of the apoplastic effector Bas4:eGFP in these experiments confirmed that the EIHM remained intact at each infection site, based on its uniform outlining of the IH and lack of spilling into the invaded rice cell.

**Figure 1.**
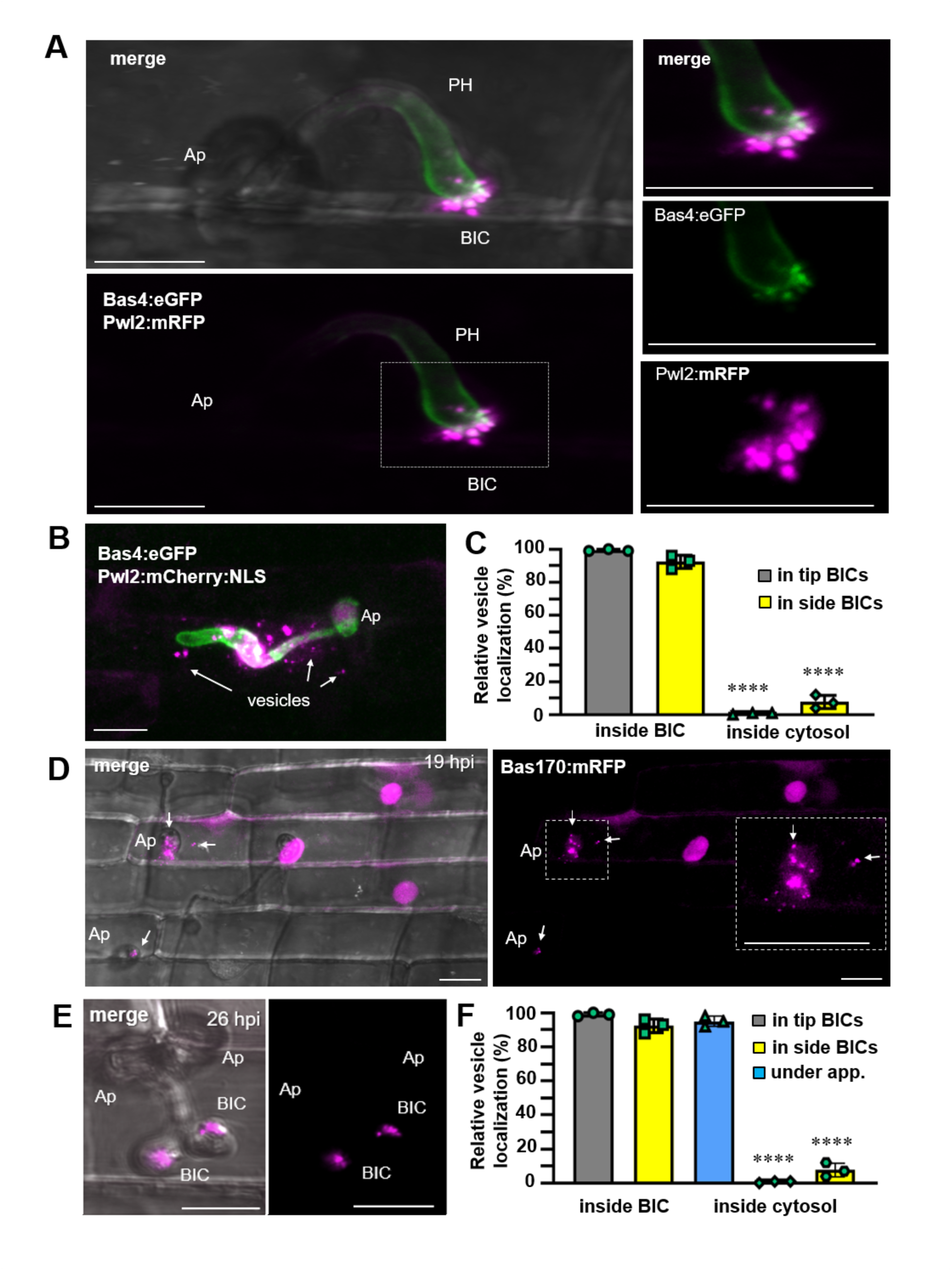
BIC vesicular organization and effector vesicles in the rice cell cytoplasm. **A.** Cytoplasmic effector-labeled vesicles in a BIC at the tip of a primary hypha (PH) of strain KV217 expressing Pwl2:mRFP and Bas4:eGFP at 17 hpi. The appressorium (Ap) and the first half of the PH are out of focus. Merged mRFP (magenta) and eGFP (green), eGFP and mRFP inset channels (right) show the base layer of apoplastic effector Bas4:eGFP and the outer layer of cytoplasmic effector Pwl2:mRFP. Occurrence of effector vesicles is independent of the cytoplasmic effector or fluorescent protein used (Supplemental Figure S1). **B.** A side-BIC occurring above a first bulbous IH cell of strain KV168 expressing Pwl2:mCherry:NLS and Bas4:eGFP at 24 hpi. Effector vesicles are visible in the host cytoplasm at a distance from the BIC (arrows). The Bas4:eGFP control shows that the EIHM surrounding the IH remains intact. See Supplemental Figure S2 for a view of the entire invaded cell showing host translocation of nuclear targeted Pwl2:mCherry:NLS. **C.** Quantification of the relative localization of cytoplasmic effector vesicles, assessed using strain KV217 from panel A. 98 vesicles carrying Pwl2:mRFP counted for each BIC developmental stage. Bar charts are based on means of all data from three biological reps (data points show the means of the individual reps); error bars indicate standard deviation. ****P<0.0001. **D.** Bas170:mRFP, here expressed by *M. oryzae* strain KV224, shows rapid accumulation in vesicles under appressoria and in plant nuclei even before a primary hypha is visible. On the right, the enlarged inset highlights vesicles near the penetration site (arrows). Left image shows merged bright field and mRFP fluorescence and right image shows mRFP fluorescence alone. **E.** Bas170:mRFP also localizes to BICs, shown here in two side-BICs resulting from adjacent appressoria penetrating the same rice cell. Experimental details same as in D. **F.** Quantification of the relative localizations of Bas170:mRFP effector vesicles, assessed using strain KV224 (D and E). 100 vesicles carrying Bas170:mRFP counted for each BIC stage and cytoplasm under appressoria (app.). Bar charts same as described in C. All images are projections of confocal optical sections. Bars = 10 µm.

We identified a novel cytoplasmic effector, Bas170 (MGG_07348.6), with amino acid sequence similarity to previously identified effector Bas83 (Mosquera et al. 2010; Supplemental Figure S3). Unlike previous cytoplasmic effectors including Pwl2, Pwl1, and Bas1, fluorescent Bas170 fusion proteins localized in vesicles in the rice cytoplasm short distances from the appressorial penetration site at a stage even before visible primary hyphal growth (Figure 1D and F). The Bas170:mRFP fusion protein naturally accumulates in the rice nucleus at this early infection stage, presumably to reprogram plant transcription essential for infection (Figure 1D and F). These results suggest rapid effector translocation from appressoria even before growth of primary IH. Bas170:mRFP also labels BIC-associated vesicles in tip-BICs and side BICs after growth of IH in the cell. The vesicular BIC-localization pattern of fluorescent Bas170 was similar to BIC-localization patterns of Pwl2-, Pwl1- and Bas1-fusion proteins (Figure 1E,F), but these latter effectors have not been observed under appressoria at the earlier time period.

### Vesicles in BICs are dynamic through time

To understand vesicle dynamics in BICs through time, we imaged BIC effector fluorescence during the morphogenic switch from tubular primary hyphae to bulbous IH using a strain co-expressing two cytoplasmic effectors, Bas1:eYFP and Pwl2:mRFP (Figure 2). The vesicle content varied over the 2-hour period when the tip-BIC was displaced from the primary hyphal tip to become a side-BIC beside the differentiating bulbous IH cell. In early stage tip-BICs, fluorescence from these two effectors often appeared to be sorted into different vesicles. For this series of experiments, Pwl2:mRFP occurred in vesicles that reached outer layers of the tip-BIC compared to Bas1:eYFP. Pwl2:mRFP appeared less abundant relative to Bas1:eYFP during the switch from tip-to side-BIC, recovering again after growth of bulbous IH (Figure 2A). Both vesicle size and content vary within BICs through time (Figure 2 B-F). Specifically, BICs at primary hyphal tips contain a larger number of smaller vesicles (Figure 2D,E). Whereas BICs on primary hyphae contain effectors in vesicles with a median diameter of 249 nm, BICs on mature bulbous hypha contain effectors in larger vesicles with a median diameter of 647 nm, ranging from ~500 nm up to above 1000 nm at later cell colonization stages (Figure 2C,D). Larger effector vesicles beside mature bulbous IH often showed colocalization of fluorescence from different effectors compared to vesicles in tip-BICs (Figure 2A, B and F).

**Figure 2.**
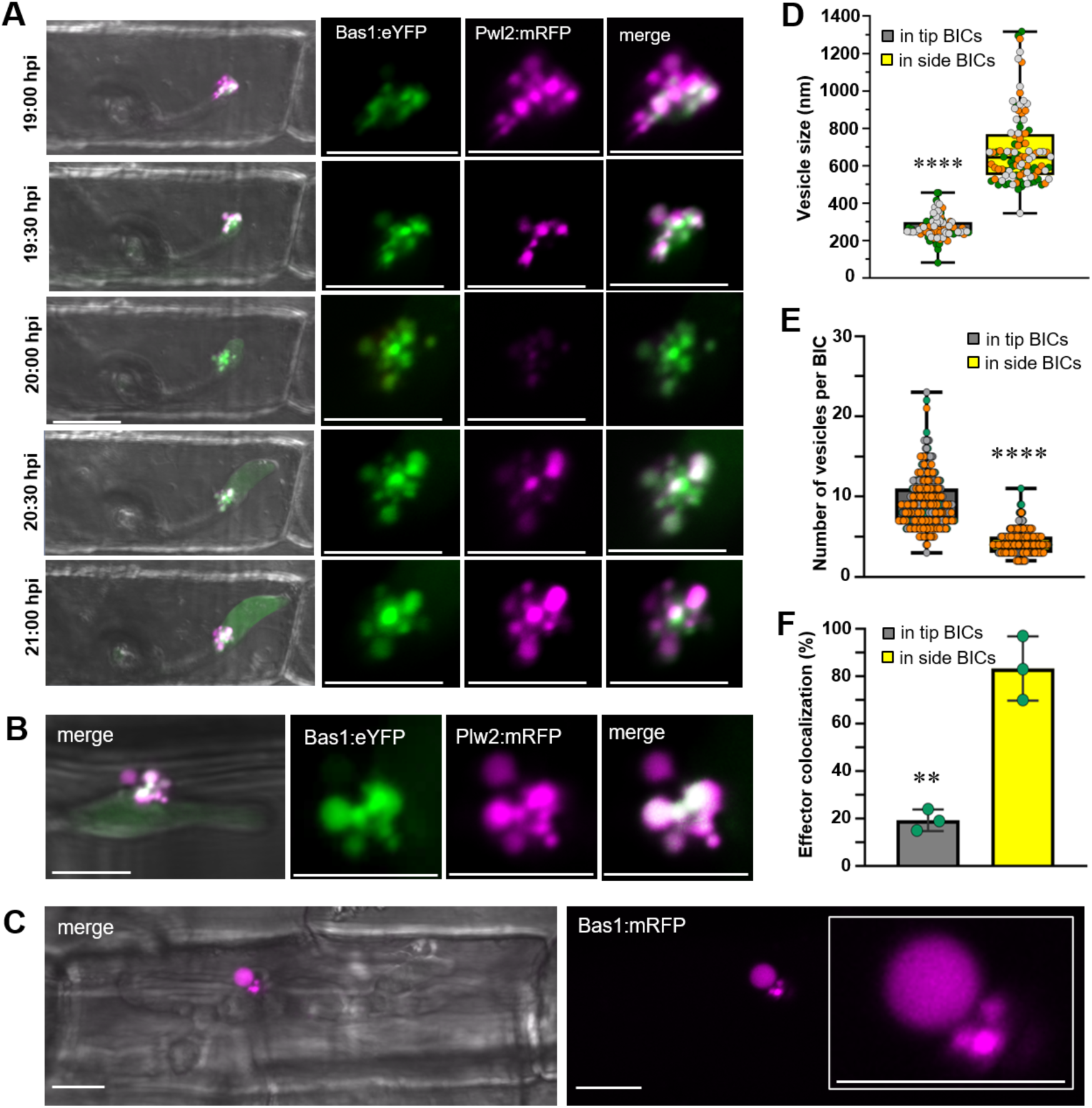
Dynamics of BICs and effector vesicles during differentiation of primary hyphae to first bulbous invasive hyphae. **A.** Two cytoplasmic effectors, Pwl2:mRFP and Bas1:eYFP (Strain KV211) were observed as a tip-BIC (top panel) becomes a side-BIC (bottom panel), imaged every 30 min from 19 to 21 hpi. Images (left to right) are merged bright field, eYFP (green) and mRFP (magneta) fluorescence at 5 time points; then enlarged views of eYFP alone, mRFP alone and merged eYFP and mRFP at same time points. Fluorescent BICs are shown as projections of confocal optical sections. Bars = 10 µm. B. Cytoplasmic effectors Pwl2:mRFP and Bas1:eYFP (Strain KV211) co-localize in larger vesicles in a side-BIC at 24 hpi. **C.** A rare extremely large effector vesicle (diameter 3.9 µm) in a late stage side-BIC formed by strain KV170 expressing Bas1:mRFP at 30 hpi. **D,E.** Box and whisker plots with individual data points comparing vesicle sizes (D) and number of vesicles (E) in tip- and side-BICs formed by strain KV211; data points of different colors represent different biological replicates. F. Bar chart showing quantification of Basl:eYFP and Pwl2:mRFP colocalization (KV211) in vesicles in tip- and side-BICs. Bar charts are based on means of all data from three biological replicates (data points show the means of the individual reps); error bars indicate standard deviation. **P 0.0015; ****P<O.OOOI. 112 vesicles measured for tip- and side-BICs in D. 100 BICs observed per replicate in E-F.

Cytoplasmic connections were observed between BICs and plant cytoplasm at the periphery of the invaded cell, illustrated in optical sections through a BIC in Supplemental Movie S1. This is consistent with a previous report of dynamic connections linking BICs and host cytoplasm underneath the appressorium (Khang et al., 2010). These cytoplasmic connections may facilitate movement of effectors from cytoplasm surrounding the BIC in the cell center to the peripheral cytoplasm, in position to rapidly move through plasmodesmata into neighboring cells.

### BIC vesicles are surrounded by plant plasma membrane

We previously reported that BICs are enriched in plant plasma membrane based on infection of transgenic rice expressing a plasma membrane marker (Giraldo et al., 2013). Here we use rice lines expressing LTi6B:GFP to confirm colocalization of plant plasma membrane and vesicles in BICs (Figure 3A,B). Individual effector vesicles were observed to be surrounded by labeled plant plasma membrane (Figure 3C), especially for vesicles in side-BICs on mature bulbous hyphae, where the vesicles reach diameters of 500 nm or larger.

**Figure 3.**
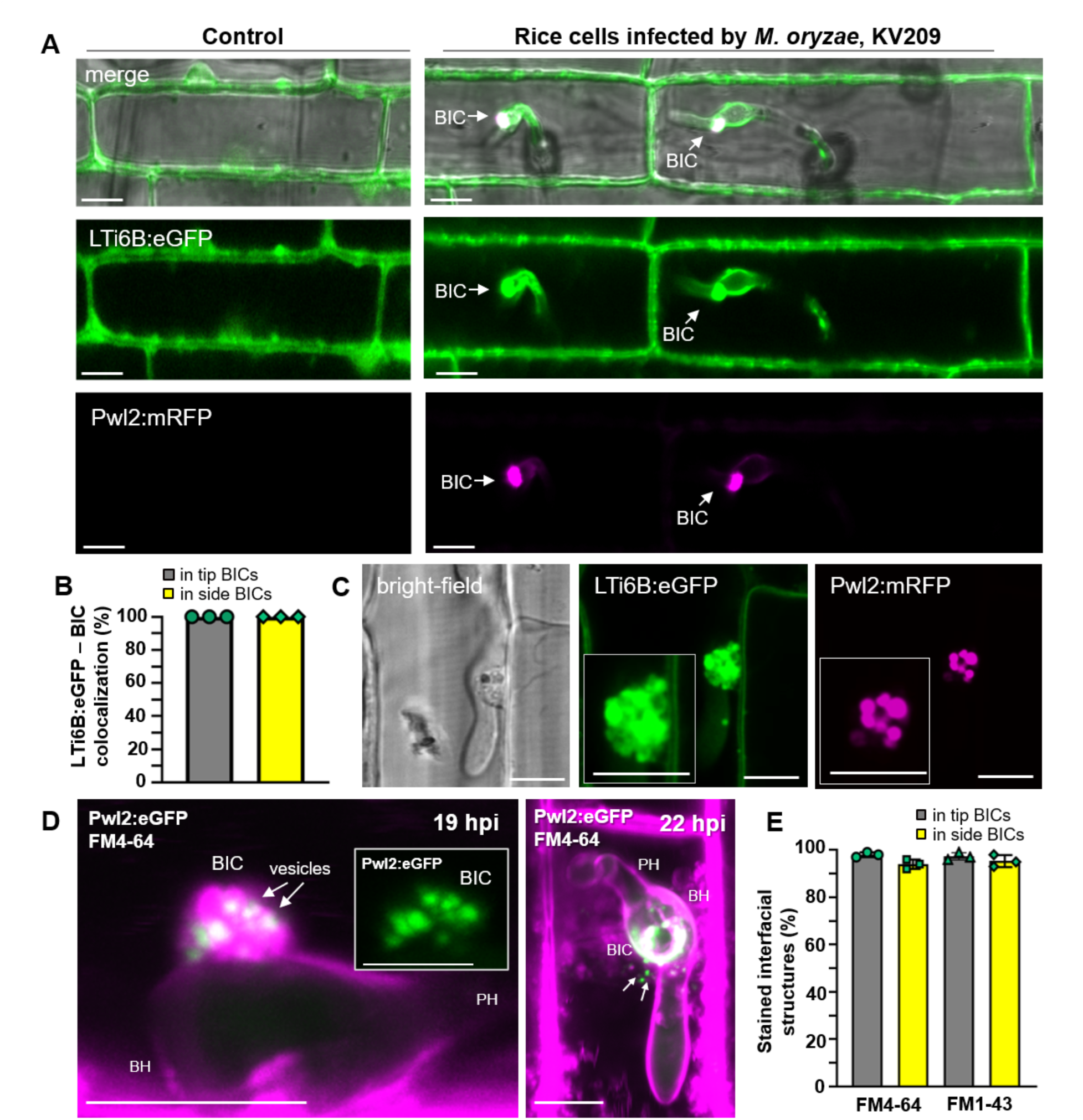
Effector vesicles are derived from plant plasma membrane. **A.** Rice expressing plasma membrane marker LTi6B:eGFP in non-infected cells (left column) and in cells infected by strain KV209 expressing Pwl2:mRFP (right column). Colocalization in a tip-BIC (left) and a side-BIC (right) confirms BICs as plant plasma membrane-rich structures. **B.** Quantification of the colocalization of LTi6B:eGFP with BICs. Bar charts are based on means of all data from three biological replicates (data points show the means of the individual reps); three times 90 infection sites observed. **C.** Localization assays of LTi6B:eGFP relative to Pwl2:mRFP in an infected rice cell at 22 hpi showing LTi6B:eGFP apparently outlining vesicles in the BIC. Insets show BIC vesicles at higher magnification. **D.** Two BICs with effector vesicles surrounded by FM4-64-stained plant membranes. Strain KV176 expressing Pwl2:eGFP shown as merged eGFP and FM4-64 fluorescence. On the left, a side-BIC at 19 hpi shows effector vesicles (eGFP fluorescence alone in the inset) immersed in FM4-64 staining. On the right, a top-down view of a different BIC at 22 hpi, image focus plane just below the BIC. Arrows indicate Pwl2:eGFP vesicles near the BIC. Images are projections of confocal optical sections. See Supplemental Movie S2 for yet another side-BIC. Note that in all three cases, lack of FM4-64 staining of the fungal septa and vacuolar membrane indicates that the EIHM was intact surrounding each IH. **E.** Quantification of BICs stained by the endocytosis tracer dyes FM4-64 or FM1-43. Bar charts are based on means of all data from three biological replicates (data points show the means of the individual reps); error bars indicate standard deviation; three times 90 infection sites observed per treatment. Bars = 10 µm.

To further confirm that BIC vesicles are associated with plant membrane, we evaluated BIC membrane composition by staining with the endocytosis tracker dyes FM4-64 and FM1-43 (Figure 3D,E). These amphiphilic styryl dyes insert in the plasma membrane where they diffuse laterally in plant and fungal membranes and are internalized into cells via endocytosis (Atkinson et al., 2002; Bolte et al., 2004). For healthy IH growing in rice cells, FM4-64 heavily stains the EIHM surrounding IH, but is excluded from IH membranes inside the EIHM compartment (Kankanala et al., 2007; Giraldo et al., 2013). This is clearly visualized by lack of lateral diffusion into IH septal membranes and lack of accumulation in IH vacuolar membranes (Giraldo et al., 2013; Kankanala et al., 2007). In these experiments, exclusion of FM4-64 or FM1-43 staining from internal IH membranes served as a control for EIHM intactness at observed infection sites (Figure 3D and Supplementary Movie S2). Quantification of FM4-64 and FM1-43 accumulation in both tip- and side-BICs (Figure 3E) confirmed that effector vesicles were immersed in the region of intense dye staining (Figure 3D; Supplementary Movie S2). Vesicles that carry effectors were infrequently observed separated from the BIC by at least 7 µm (Figure 3D left, right; Supplemental Movie S2). Apparent vesicular bursting was also observed, which suggests that the vesicles are unstable, presumably releasing the effector contents into the host cytoplasm. Taken together, results from labeling of rice plasma membrane with both LTi6B:GFP membrane fusion protein and endocytosis tracer dyes are consistent with the formation of BIC-associated effector vesicles through plant endocytosis.

### Cytoplasmic effector Bas83 associates with rice membrane vesicles concentrated around BICs

Since it is likely that specific blast effectors play roles in vesicle formation and effector translocation, we were particularly interested in effectors that appear to associate with membranes in host cells. The *BAS83* (MGG_08506.6) gene is unique to *M. oryzae*, and it is 36-fold upregulated during biotrophic invasion (Mosquera et al., 2009). Bas83:mRFP is translocated into invaded host cells, but it does not precisely colocalize with other BIC-localized cytoplasmic effectors Pwl2:eGFP and Pwl1:eGFP (Figure 4A; Supplemental Figure S4). In contrast, Bas83:mRFP seems to label the EIHM surrounding the BIC-associated cells, the primary hyphae and first bulbous IH, and it labels bubbles, or vesicles, near to BICs and BIC-associated cells (Figure 4A-C). Figure 4A shows that Bas83:mRFP-labeled vesicles do not contain the cytoplasmic effector Pwl2:eGFP, indicating they are distinct from effector vesicles in BICs. Bas83:mRFP is also associated with vesicles observed under appressoria (Figure 4C; Supplemental Figure S4). However, Bas83:mRFP fails to label membrane surrounding subsequently formed bulbous IH that are associated with apoplastic effectors and not with cytoplasmic effectors (see 2^nd^ bulbous IH in Figure 4C). We hypothesize that Bas83 is involved in recruiting new plant plasma membrane to form the EIHM and BICs during biotrophic development. However, repeated attempts in two different laboratories (264 purified transformants, 4 methods and 5 experiments; KSU and Univ. of Exeter) failed to produce knock-out mutants for further functional analysis (Supplemental Figure S5).

**Figure 4.**
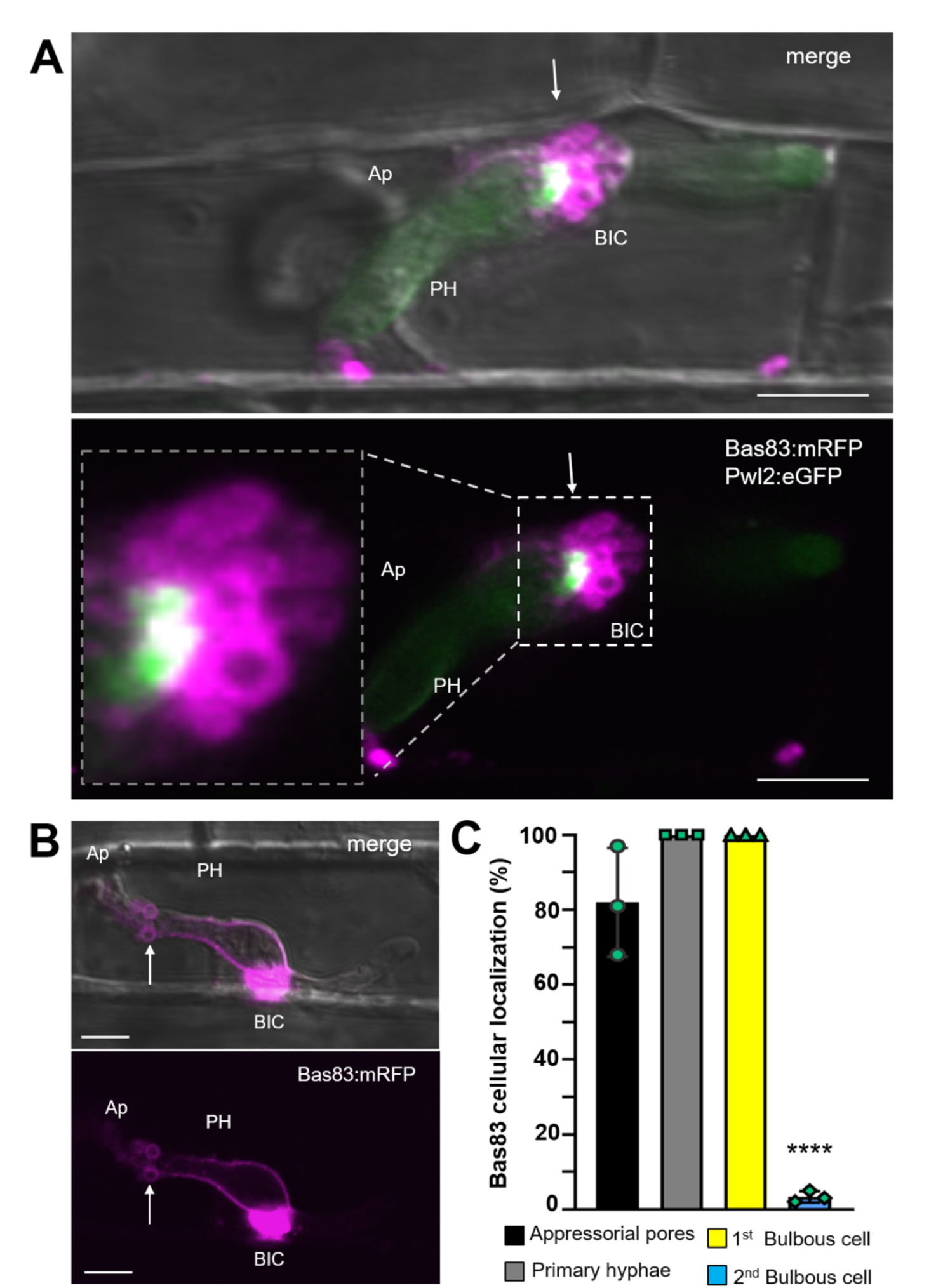
Bas83 is a novel effector associated with membrane vesicles surrounding BICs and BIC-associated cells. **A.** Bas83:mRFP binds apparent membrane vesicles surrounding a side-BIC (white arrow) labeled with Pwl2:eGFP (strain KV222). **B.** Bas83:mRFP (KV220) binds to bubble-like structures near a primary hypha (white arrow) as well as labeling a BIC and outlining BIC-associated cells. Images (merged bright field, eGFP and mRFP fluorescence) are projections of confocal optical sections. See additional images in Supplemental Figure S4. Bars = 10 µm. **C.** Quantification of the cellular localization of Bas83:mRFP during biotrophic invasion of *M. oryzae* KV220. Bar charts are based on means of all data from three biological replicates (data points show the means of the individual reps); error bars indicate standard deviation. ****P<0.0001; 100 infection sites observed per experiment.

### BICs contain plant actin

Although it is now clear that actin is dispensable for CME vesicle formation at the plasma membrane in plant cells, actin filaments likely play a role in short-distance movement of internalized vesicles (Narasimham et al., 2020; Ruan et. al., 2021: Schuh, 2011). Therefore, we localized plant actin during biotrophic invasion using rice lines expressing the actin marker LifeAct:eGFP. Infection by *M. oryzae* strains expressing chimeric protein Pwl2:mRFP showed localization of LifeAct:eGFP at BICs (Figure 5A). Actin was abundant in all imaged tip- and side-BICs (Figure 5B). Likewise, colocalization of fluorescent Phalloidin conjugates and fluorescently-labeled effector confirmed that BICs are rich in plant actin filaments (Figure 5C). We hypothesize that actin filaments are responsible for the transport of effector vesicles in the rice cell cytoplasm.

**Figure 5.**
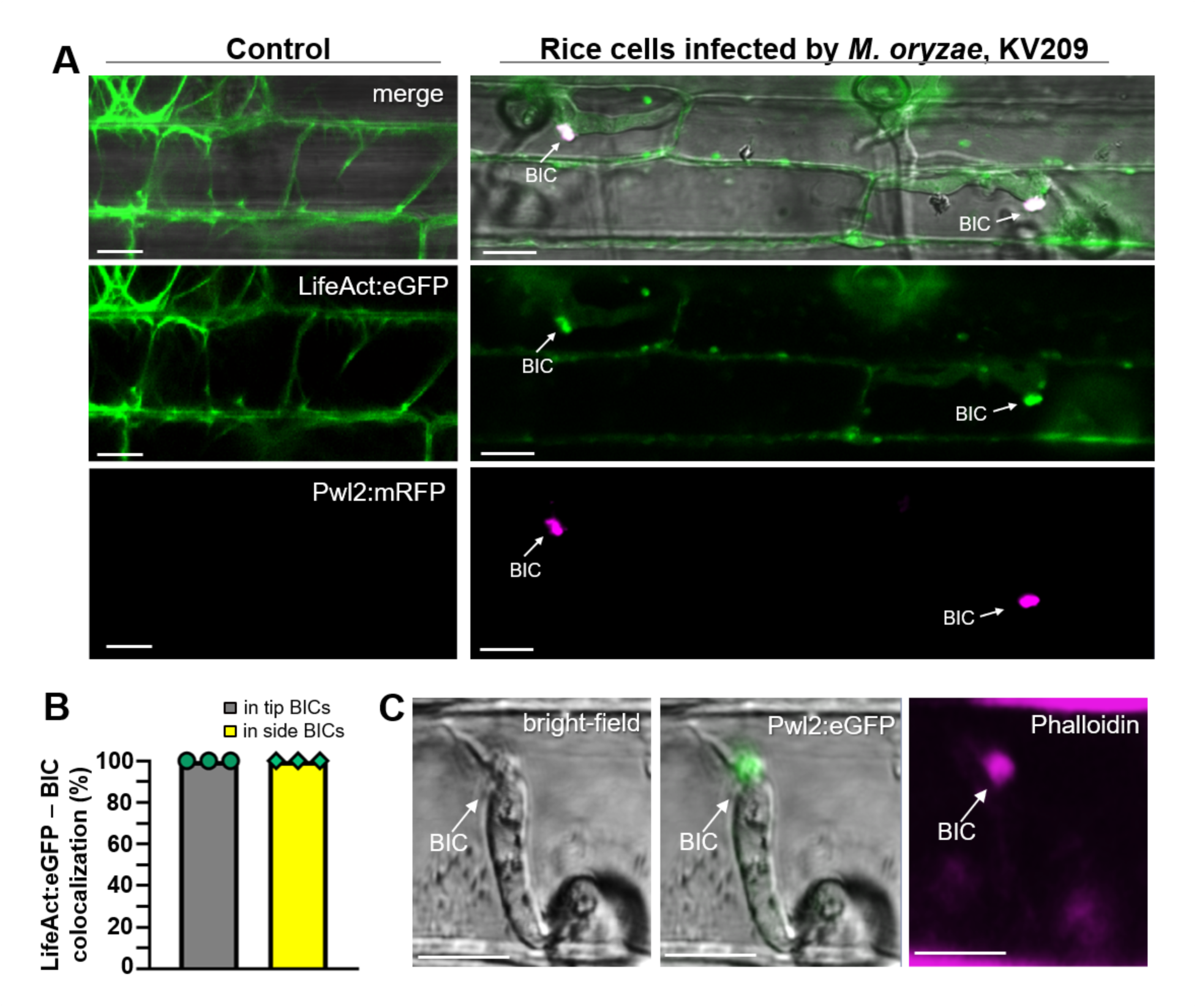
BICs contain plant actin. **A.** Rice LifeAct:eGFP localization during biotrophic invasion by *M. oryzae*. A rice line expressing LifeAct:eGFP in non-infected cells (left column) and cells infected by *M. oryzae* strain KV209 expressing Pwl2:mRFP in BICs (right column). Colocalization shows BICs are rich in plant actin. **B.** Quantification of the colocalization of LifeAct:eGFP with both tip- and side-BICs. Bar charts are based on means of all data from three biological replicates (data points show the means of the individual reps); error bars indicate standard deviation; three times 100 infection sites observed. **C.** Rhodamine phalloidin conjugate (red fluorescence staining actin, shown as magneta) colocalizes with Pwl2:eGFP (green) in rice cells infected by strain KV176, confirming the actin-rich structure of the BIC.

### BICs consistently contain plant clathrin

Clathrin-mediated endocytosis (CME) is the major endocytosis mechanism in plants, with some contribution by Clathrin-independent endocytosis (CIE) (Chen et al., 2011; Ewers and Helenius, 2011; Fan et al., 2015; Narasimhan et al., 2020). To assess localization of specific endocytosis markers in BICs, we generated transgenic rice lines expressing a fluorescent marker for either CME or CIE. For CME, we generated a C-terminal translational fusion of eGFP with the rice clathrin light chain-1 (OsCLC1), which is a component of the clathrin coat together with the clathrin heavy chain. Rice lines expressing OsCLC1:eGFP show typical clathrin foci (Konopka et al., 2008; Narasimhan et al., 2020) in uninfected rice cells (Figure 6). The marker for CIE was a C-terminal translational fusion of eGFP to Flotillin-1 (Flot1), which is a component of lipid rafts (Otto and Nichols, 2011; Li et al., 2012). Rice lines with the CME and CIE markers were used for leaf sheath assays with an *M. oryzae* strain expressing Pwl2:mRFP to label BICs. In addition to typical clathrin foci around the cell periphery, all tip-BICs and side-BICs contain fluorescence from OsCLC1:eGFP co-localizing with Pwl2:mRFP (Figure 6A,B and E). Indeed, clathrin fluorescence was observed co-localizing with Pwl2:mRFP or Bas1:mRFP effector fluorescence in BIC vesicles in both tip and side-BICs (Figure 6A-C; Supplemental Figure S6; Supplemental Movie S3). In contrast, very few (~4%) tip-BICs colocalized with Flot1:eGFP. Some colocalization of Flot1:eGFP and Pwl2:mRFP occurred in side-BICs besides mature bulbous hyphae (Figure 6D,E). Colocalization results from several independent experiments were consistent with a major role for CME, but not for CIE, in the internalization of cytoplasmic effectors.

**Figure 6.**
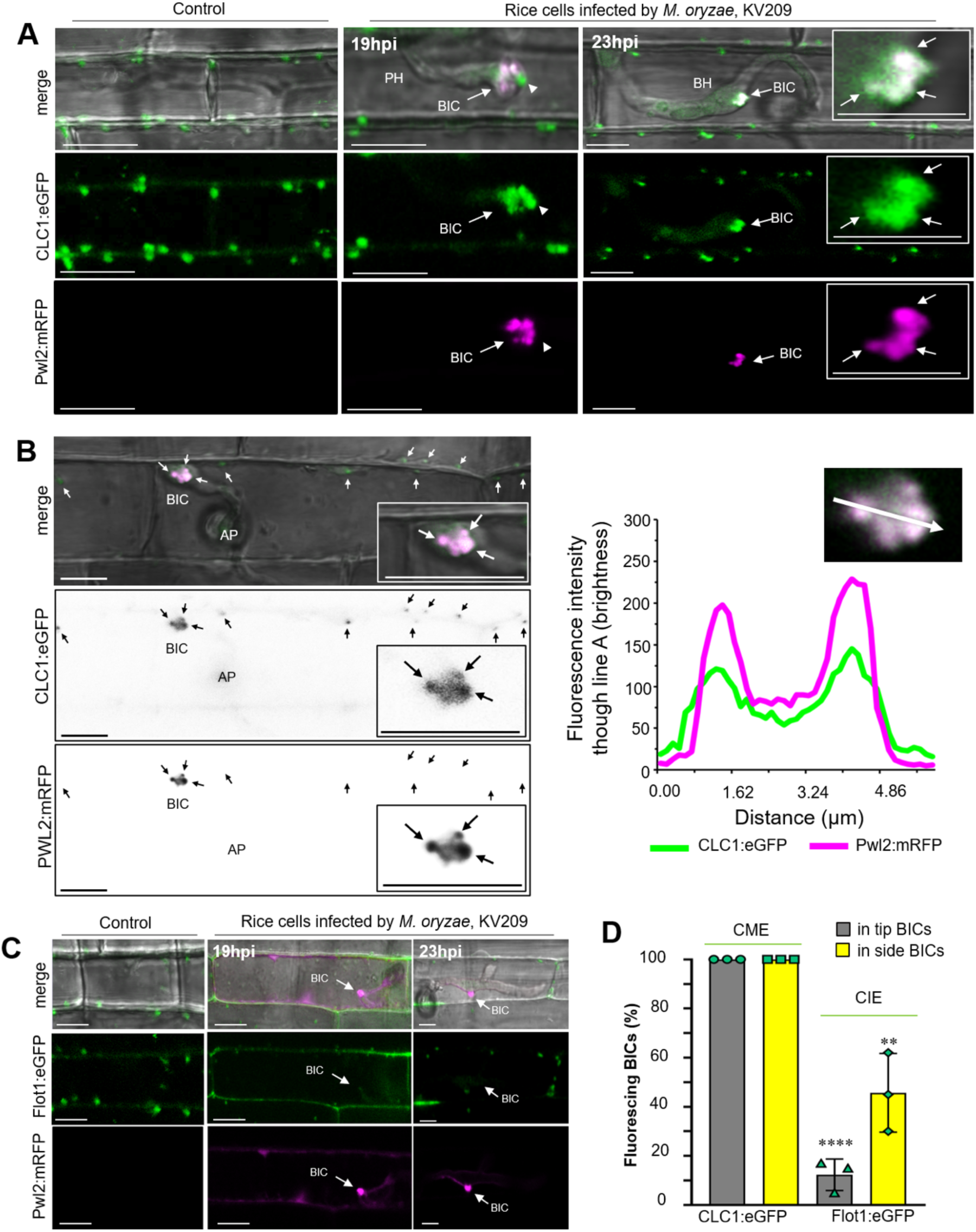
BICs uuifonuly contain clathrin, but not in flotillinl. **A.** Rice leaf sheaths expressing CME marker OsCLCl:eGFP were inoculated with strain KV209 expressing Pwl2:mRFP. Shown from left to right are a non-infected rice sheath cell; an infected cell containing a primary hypha (PH) with a tip-BIC; and an infected cell containing a bulbous IH (BH) with a side-BIC. Arrows indicate BICs. Note a vesicle visualized with OsCLC1:eGFP in the tip-BIC (arrowhead) that lacks Pwl2:mRFP, which is consistent with results in Figure 2A that fluorescent effectors are not always distributed uniformly in vesicles. (Bars=10µm.) **B.** The fluorescence pattern of OsCLC1:eGFP (green) corresponds to the Pwl2:mRFP vesicular pattern (magenta), as shown by a fluorescence intensity linescan along the path of the white arrow. The merge image shows the brightfield view together with red (magenta) and green fluorescence. Individual black and white inverse images of CLC1:eGFP and Pwl2:mRFP. Note that this image has been extracted from Supplemental Movie S3, which shows OsCLC1-labeled effector vesicles in time lapse during tip-BIC to side-BIC differentiation. **C.** Rice leaf sheaths expressing OsFlot1:eGFP (CIE marker) were inoculated with KV209 expressing Pwl2:mRFP. From left to right, shown are OsFlot1:eGFP fluorescence in a non-infected sheath cell; lack of OsFlot1:eGFP fluorescence in a tip-BIC; and lack of OsFlot1:eGFP fluorescence in a side-BIC. Arrows indicate BICs. Bars=10µm. **D.** Quantification of the localization of OsCLC1:eGFP or OsFlot1:eGFP at both BIC developmental stages. Bar charts are based on means of all data from three biological replicates (data points show the means of the individual reps); error bars indicate standard deviation; three times 97 BICs observed each treatment. ****P<0.0001; **P=0.002. CME, clathrin-mediated endocytosis; CIE, clathrin-independent endocytosis.

### Silencing of rice endocytosis machinery blocks *M. oryzae* infection and impacts BIC structure

We assessed the impact of inhibition of CME and CIE on BICs and effector vesicles using the brome mosaic virus system (Ding et al., 2006; Ding et al., 2007) for silencing of genes for CME and CIE components previously characterized in *Arabidopsis*. Specifically, we silenced two rice CME genes, Adaptor protein complex-2 subunit a2 (*AP-2*a) and clathrin heavy chain-1 (*CHC1*) (Di Rubbo et al., 2013; Larsson et al., 2017) as well as the CIE gene *Flotillin-1* (*Flot1*) (Li et al., 2012). Silencing of both CME components *AP-2*a and *CHC1* resulted in severe reduction in pathogenicity in whole plant spray inoculation assays (Figure 7A,B). However, as expected, silencing of these genes caused stunting and decreased plant health (Supplemental Figure S7), and levels of pathogenicity in blast disease are known to be decreased on unhealthy rice plants. More significant is the phenotype of pathogen blockage at individual infection sites. Silencing of either CME component RNAi*AP-2*a or RNAi*CHC1* led to a distinctive swollen BIC phenotype, with enlarged, abnormally-shaped BICs that lack effector vesicles (Figure 7C-F). The swollen BIC phenotype would be consistent with effectors still being secreted by the fungus, but remaining trapped within the EIHM due to blocking of CME. By contrast, silencing of the CIE marker gene *Flot1* showed less severe stunting (Supplemental Figure S7), less of an impact on pathogenicity (Figure 7A,B), and minor impact, if any, on BIC structure (Figure 7C-F). For example, compare the 28 hours post-inoculation (hpi) images of BICs in control rice plants with the RNAi*Flot1-*silenced and the RNAi*AP-2*a*-*silenced rice plants in Figure 7C, and quantification of swollen BIC events in Figure 7F.

**Figure 7.**
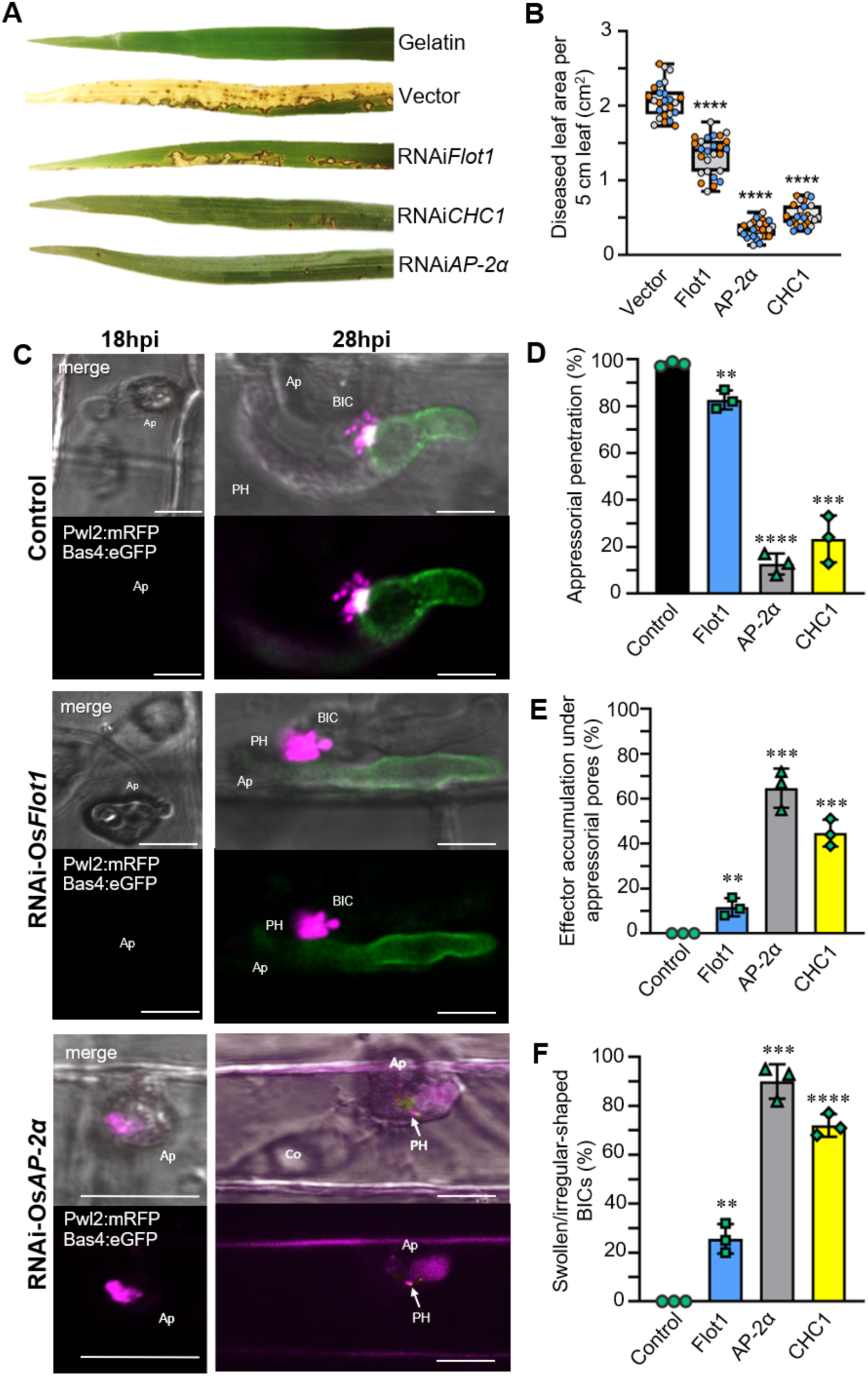
Silencing of rice endocytic machinery suggests roles in pathogenicity and fungal effector translocation. **A.** Severity of disease caused by *M. oryzae* strain Guy11 in rice leaves, cultivar IR64, undergoing silencing of rice endocytic genes. Whereas silencing of *Flot1* gene showed minor effects on pathogenesis, the silencing of the key clathrin-mediated endocytosis genes, *AP-2*a and CHC1 resulted in a significant reduction in pathogenicity in whole plant spray inoculation assays. **B.** Quantification of diseased area in 5cm leaf segments in rice expressing RNAi*AP-2*a, RNAi*CHC1*, RNAi*Flot1* or empty vector infected by strain Guy11. Box and whisker plots with individual data points are shown; data points of different colors represent different biological replicates. (P<0.0001 for all treatments; n = 9 rice plants per replication). **C.** Effects of silencing rice endocytosis genes on the localization of Pwl2:mRFP and Bas4:eGFP during biotrophic invasion. Wild type IR64 rice and rice expressing RNAi*Flot1* or RNAi*AP-2*a were infected by KV217 expressing Pwl2:mRFP and Bas4:eGFP at 26 hpi. Note the accumulation of Pwl2:mRFP under the appressorium (Ap) and the pale Bas4:eGFP fluorescence associated with the short primary hypha (PH) in the RNAi*AP2*a rice compared to the control and RNAi*Flot1* rice. Images in C (merged bright field, eGFP and mRFP fluorescence) are shown as projections of confocal optical sections. Bars = 10 µm. **D.-F.** Quantification of appressorial penetration (D), effector accumulation under appressoria (E), and swollen irregular BICs (F) in rice expressing RNAi*Flot1*, RNAi*AP-2*a, RNAi*CHC1* or the empty vector during infection by strain KV209 expressing Pwl2:mRFP (~28 hpi). Bar charts are based on means of all data from three biological replicates (data points show the means of the individual reps); error bars indicate standard deviation. (D: **P=0.0031, ***P=0.0002, ****P<0.0001; E: **P=0.0075, ***P=0.0002; F: **P=0.0018, ****<0.0001; three times 100 infection sites counted per treatment).

In addition to swollen BICs at sites with primary hyphae, silencing of both CME genes decreased appressorial penetration and induced the accumulation of the cytoplasmic effector Pwl2:mRFP under failed penetration sites, where it is not normally observed (Figure 7 C-18hpi,D,E). In contrast, silencing of *Flot1* had only minor effects on appressorium penetration and on accumulation of Pwl2:mRFP under failed appressoria. Similar to our finding of early accumulation of Bas170 under appressoria before penetration, these results again suggest that effector uptake begins before host penetration. These results also suggest that CME plays a role in effector translocation at the appressorial penetration stage as well as from BICs.

### Chemical inhibition of rice endocytosis machinery blocks *M. oryzae* infection

Pharmacological approaches have impacted studies in diverse biological systems (Robinson et al., 2008; Hicks and Raikhel, 2010; Grassart et al., 2014; Fan et al., 2015). To further assess potential impact of the plant endocytic machinery on cytoplasmic effector translocation, we tested a series of chemicals that are reported to inhibit plant CME or CIE (Table 1). Compared with the non-treated control (Figure 8A), treatment with CME inhibitor Endosidin9-17 (ES9-17), which directly inhibits clathrin heavy chain function in both *Arabidopsis* and human cells (Dejonghe, et al., 2019), led to abnormally-shaped swollen BICs that lack distinct effector vesicles, resembling those seen with silencing of CME components CHC-1 and *AP-2*a (Figure 8B). Treatment with cantharidin, which is reported to inhibit signaling associated with CME, also produced swollen BICs lacking effector vesicles (Figure 8C). Fluorescence intensity linescans confirm that ES9-17 and cantharidin had little, if any, impact on localization of apoplastic effector Bas4:eGFP in the base of the BIC (Figure 8).

**Figure 8.**
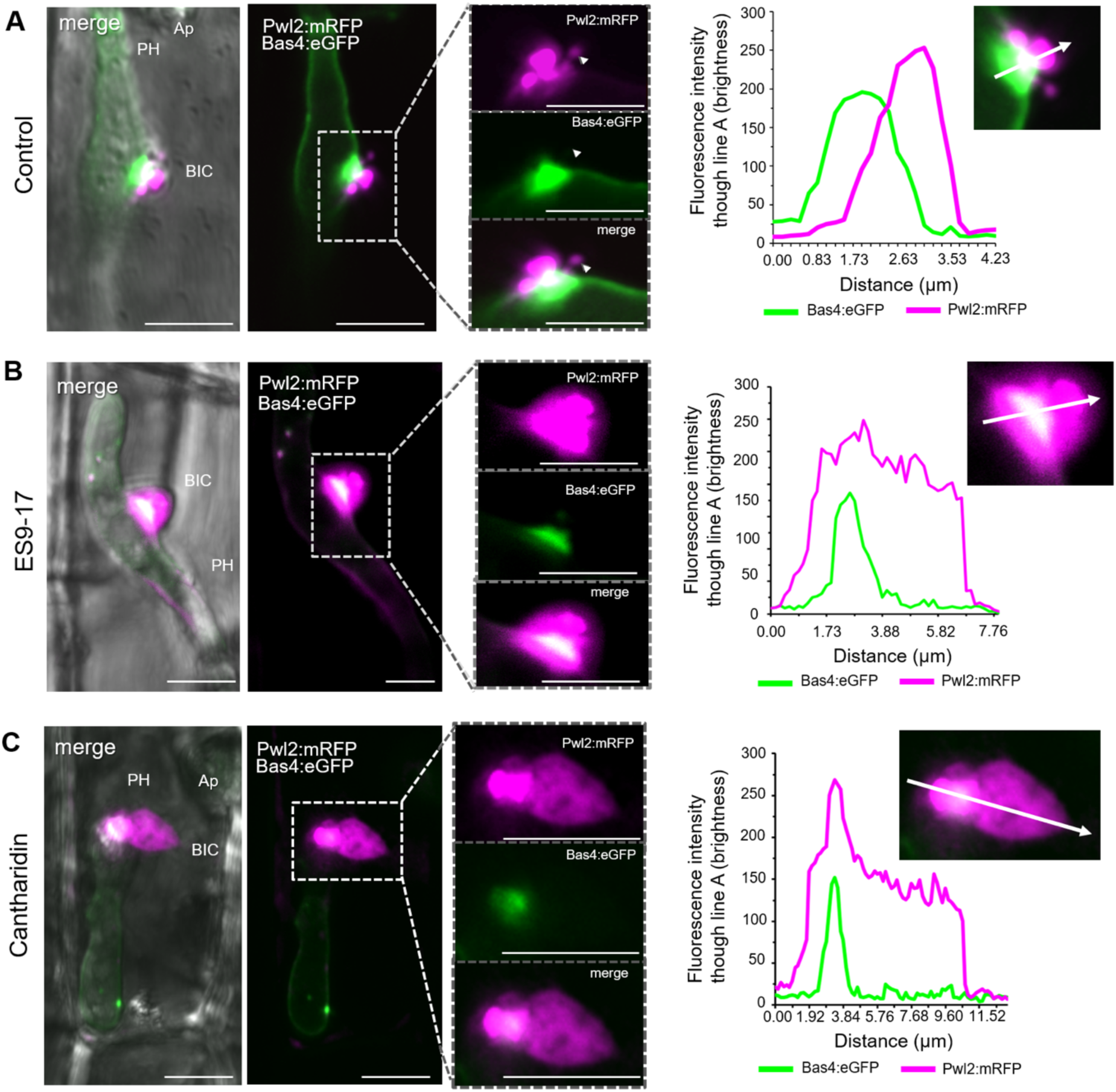
Treatment with CME inhibitor ES9-17 and CME signaling inhibitor cantharidin impacts the localization of cytoplasmic effector Pwl2, but not apoplastic effector Bas4 in BICs. All panels show side-BICs formed during invasion of rice by strain KV217 expressing Pwl2:mRFP and Bas4:eGFP at 26 hpi. **A.** Apoplastic effector Bas4:eGFP identifies an inner base layer of the BIC relative to cytoplasmic Pwl2:mRFP, as shown by fluorescence intensity linescans for GFP (green) and mRFP (magenta) along the path of the white arrow. **B.** After treatment with clathrin heavy chain inhibitor ES9-17, localization of Bas4:eGFP in the BIC base appears unaffected, compared to the non-vesicular swollen localization of Pwl2:mRFP. The white arrows in the inset show the paths for fluorescence intensity distribution linescans for GFP (green) and mRFP (magenta). **C.** After treatment with CME inhibitor cantharidin, localization of Bas4:eGFP in the BIC base appears unaffected, compared to the non-vesicular swollen localization of Pwl2:mRFP, as shown by fluorescence intensity linescans for GFP (green) and mRFP (magenta) along the path of the white arrow. Note linescan distances for Pwl2:mRFP are 2.5 to 3-fold greater after inhibitor treatments. From left to right, images show merged bright field, eGFP and mRFP fluorescence; merged eGFP and mRFP fluorescence; insets with enlarged mRFP, eGFP and merge views; and intensity linescans along the paths of the white arrows. Images are projections of confocal optical sections. Appressorium (Ap); Primary hypha (PH). Bars10 11m.

**Table 1.**
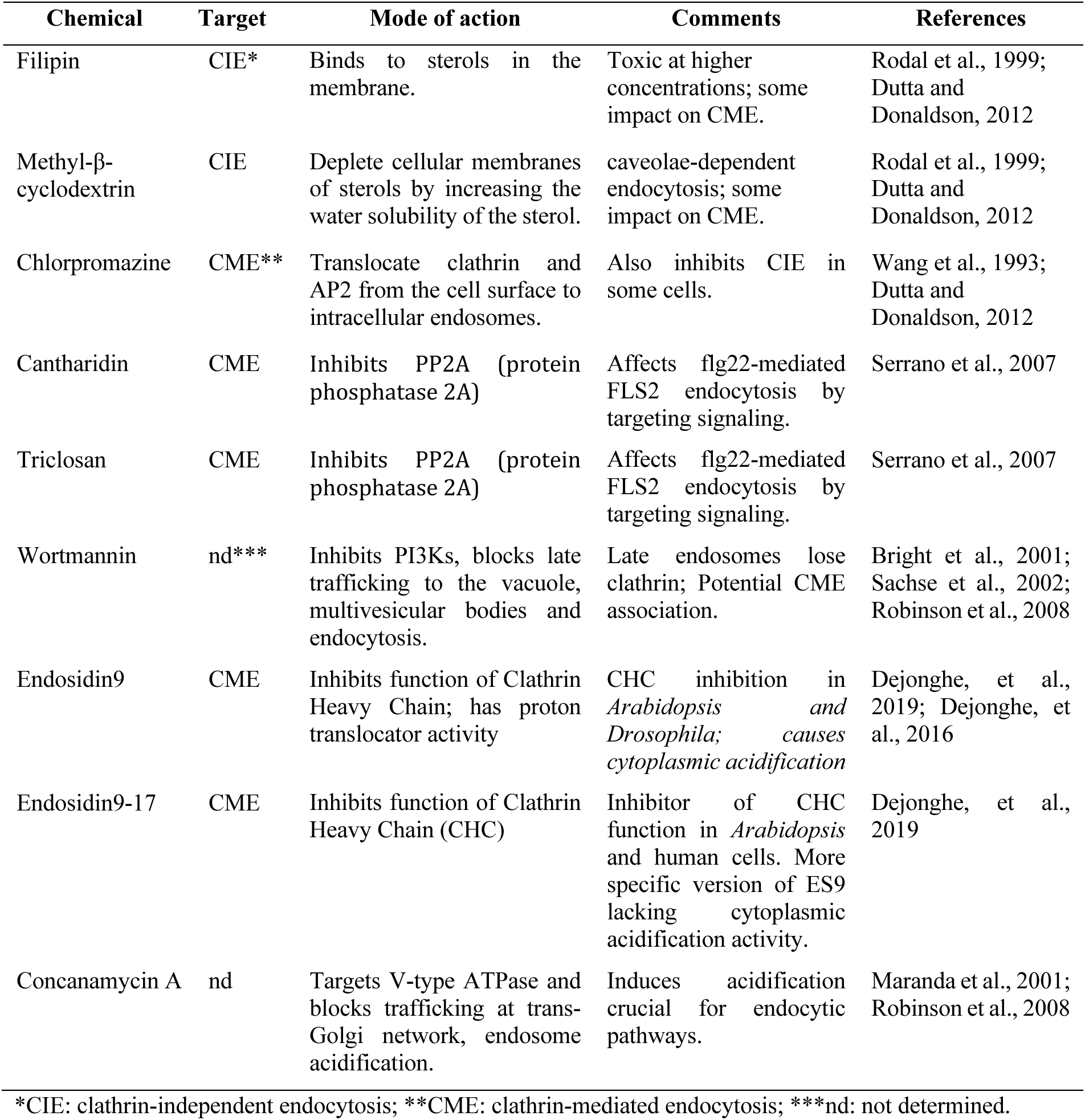
Chemical inhibitors of endocytosis

| Chemical | Target | Mode of action | Comments | References |
| --- | --- | --- | --- | --- |
| Filipin | CIE* | Binds to sterols in the membrane. | Toxic at higher concentrations; some impact on CME. | Rodal et al., 1999; Dutta and Donaldson, 2012 |
| Methyl-β-cyclodextrin | CIE | Deplete cellular membranes of sterols by increasing the water solubility of the sterol. | caveolae-dependent endocytosis; some impact on CME. | Rodal et al., 1999; Dutta and Donaldson, 2012 |
| Chlorpromazine | CME** | Translocate clathrin and AP2 from the cell surface to intracellular endosomes. | Also inhibits CIE in some cells. | Wang et al., 1993; Dutta and Donaldson, 2012 |
| Cantharidin | CME | Inhibits PP2A (protein phosphatase 2A) | Affects flg22-mediated FLS2 endocytosis by targeting signaling. | Serrano et al., 2007 |
| Triclosan | CME | Inhibits PP2A (protein phosphatase 2A) | Affects flg22-mediated FLS2 endocytosis by targeting signaling. | Serrano et al., 2007 |
| Wortmannin | nd*** | Inhibits PI3Ks, blocks late trafficking to the vacuole, multivesicular bodies and endocytosis. | Late endosomes lose clathrin; Potential CME association. | Bright et al., 2001; Sachse et al., 2002; Robinson et al., 2008 |
| Endosidin9 | CME | Inhibits function of Clathrin Heavy Chain; has proton translocator activity | CHC inhibition in *Arabidopsis* and *Drosophila*; causes cytoplasmic acidification | Dejonghe, et al., 2019; Dejonghe, et al., 2016 |
| Endosidin9-17 | CME | Inhibits function of Clathrin Heavy Chain (CHC) | Inhibitor of CHC function in *Arabidopsis* and human cells. More specific version of ES9 lacking cytoplasmic acidification activity. | Dejonghe, et al., 2019 |
| Concanamycin A | nd | Targets V-type ATPase and blocks trafficking at trans-Golgi network, endosome acidification. | Induces acidification crucial for endocytic pathways. | Maranda et al., 2001; Robinson et al., 2008 |

\*CIE: clathrin-independent endocytosis; \*\*CME: clathrin-mediated endocytosis; \*\*\*nd: not determined.

We quantified Pwl2:mRFP localization patterns during infection of rice with no treatment, after treatment with five chemicals reported to impact CME inhibitors, two chemicals reported to impact CIE, and 2 chemicals with less defined roles in inhibiting endocytosis (Figure 9). The five CME inhibitors included ES9-17 and cantharidin (Figure 8) along with additional reported CME inhibitors, Endosidin9 (ES9), chlorpromazine and triclosan (Table 1). Compared to the untreated control, each of these five CME-associated inhibitors showed a high proportion of swollen, irregular-shaped cytoplasmic effector fluorescence patterns in BICs (Figure 9A-C). By contrast, inhibition of CIE with filipin and methyl-β-cyclodextrin resulted in minor to no effect on BIC effector fluorescence patterns (Figure 9A-C), which was consistent with results from silencing of CIE (Figure 8). Two chemical inhibitors with potential CME association (Table 1) show different impacts on BIC effector fluorescence patterns. Wortmannin, which inhibits phosphatidylinositol 3-kinase (PI 3-kinase) and is implicated in late endosomal trafficking (Robinson et al., 2008) gave the swollen BIC phenotype, while concanamycin A, which inhibits vacuolar-type ATPase and induces vacuolar acidification (Robinson et al., 2008), had little impact on BIC structure (Figure 9A-C).

**Figure 9.**
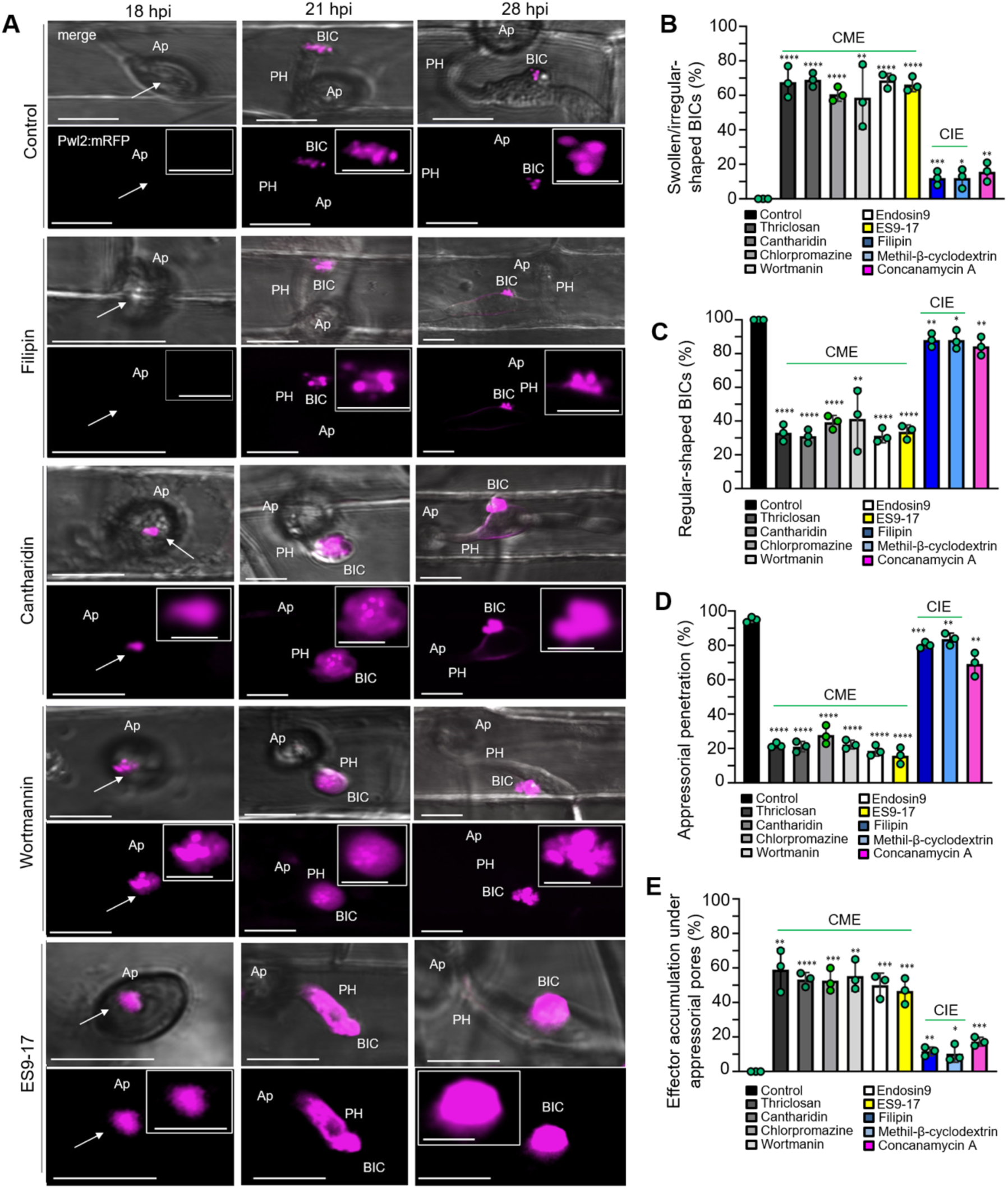
Chemical inhibition of rice CME, but not of CIE, has a major impact on cytoplasmic effector localization in BICs and under appressoria. The chemical inhibitors used are described in Table 1. **A.** Effects of chemical inhibition of clathrin-mediated endocytosis (CME) and clathrin-independent endocytosis (CIE) in the rice sheath assay with strain KV209 expressing Pwl2:mRFP at 18, 21 and 28 hpi. Whereas inhibition of CIE showed minor, although significant, effects on BIC functionality and effector uptake, the chemical inhibition of CME resulted in a significant accumulation of effector fluorescence under appressoria (Ap, white arrows at 18 hpi) as well as swollen BICs on primary hyphae (PH, 21 hpi) and bulbous IH (28hpi). Images in A (merged bright field and mRFP fluorescence) are projections of confocal optical sections. Insets show close-up views of the effector fluorescence. Bars = 10 µm and bars in the insert boxes = 5 µm. **B-E.** Quantification of swollen, irregular-shaped BICs (B), regular-shaped BICs (C), appressorial penetration (D), and effector accumulation under appressoria (E), in rice undergoing chemical treatment and infection by strain KV209 expressing Pwl2:mRFP at 28 hpi. Bar charts are based on means of all data from three biological replicates (data points show the means of the individual reps); error bars indicate standard deviation. Three times 100 infection sites observed per treatment. B: *P=0.0202(MβC), **P=0.0065(Filipin), **P=0.0079 (Con.A), **P=0.005(Wort.), ****P<0.0001; C: *P=0.02(MβC), **P=0.0065(Filipin), **P=0.0075(Con.A), **P=0.005(Wort.), ****P<0.0001; D: P=0.0068(MβC), **P=0.0004(Filipin), **P=0.0032 (Con.A), ****P<0.0001; E: *P=0.0222(MβC), **P=0.0013(Filipin), ***P=0.0003(Con.A), ***P=0.0002(ES9), ***P=0.0003(ES9-17), **P=0.0004(Wort.), **P=0.0011(Tricl.), ***P=0.0002(Chlorpr.), ****P<0.0001(Canth.).

Moreover, treatment with inhibitors associated with CME, triclosan, cantharidin, chlorpromazine, ES9, and ES9-17, but not with inhibitors associated with CIE, filipin and methyl-β-cyclodextrin, resulted in reduced appressorial penetration and induced abnormal accumulation of the cytoplasmic effector Pwl2:mRFP under appressorial penetration sites where it is not normally observed (Figure 9A - compare 18 hpi CIE and CME inhibitors; Figure 9D,E). Wortmannin, which produces swollen BICs, also impacted appressorial function and resulted in abnormal Pwl2:mRFP accumulation under appressoria (Figure 9D,E). Concanamycin A produced none of the CME-associated phenotypes (Figure 9). Therefore, chemical inhibition of endocytosis in rice cells undergoing *M. oryzae* infection mimics the results obtained with VIGS, further supporting a role for CME in effector uptake into rice cells. These CME inhibition results support our findings with Bas170:mRFP (Figure 1D-F) in suggesting that effector uptake begins before or at the point that host penetration occurs.

## DISCUSSION

Our results, when taken together, provide strong evidence that cytoplasmic effector translocation is BIC-localized and mediated by clathrin-mediated endocytosis (CME). We report that BICs contain multiple hallmarks of active CME. Specifically, BICs uniformly show focal accumulation of fluorescently-labeled Clathrin Light Chain-1 (CLC1:eGFP), a key subunit of the clathrin coat in CME (Figure 6). Indeed, CLC1:eGFP fluorescence has frequently been observed in a pattern co-localizing with individual cytoplasmic effector vesicles (Figure 6A and B; Supplemental Figure S6; Supplemental Movie S3). Fluorescent rice plasma membrane fusion protein LTi6B:eGFP colocalizes with effector vesicles (Figure 3C, Supplemental Movie S2). Results with endocytosis tracer dyes FM4-64 or FM1-43 served both as a control that the EIHM was intact and excluding the dye from reaching fungal membranes at imaged infection sites (Kananala et al., 2007; Giraldo et al., 2013), and showed effector vesicles emersed in stained plant membranes (Figure 3D and E). Functional analyses via virus induced gene silencing of two rice CME components, Clathrin Heavy Chain-1 (CHC-1) and Adaptor Protein Complex subunit 2α (AP-2α) decreased pathogenicity and resulted in a ‘swollen BIC’ phenotype in which cytoplasmic effectors appear retained inside an enlarged BIC compartment lacking distinct vesicles (Figure 7). In contrast, parallel experiments showed that fluorescently-labeled clathrin independent endocytosis (CIE) component Flotillin-1 (Flot1:eGFP) rarely localized at BICs, and silencing of Flotillin-1 had minimal effects on BIC vesicular structure (Figure 7). Additionally, treatment with 6 different pharmacological agents associated with CME, including the direct inhibitor of clathrin heavy chain function Endosidin9-17, resulted in the swollen BIC phenotype. Parallel experiments with pharmacological inhibitors associated with CIE show little or no impact on BIC vesicular structure and pathogenicity (Figure 8 and 9). Taken together, our results therefore provide evidence that BICs include a focused region of plant clathrin-mediated endocytosis for internalizing cytoplasmic effectors inside living plant cells (Figure 10).

**Figure 10.**
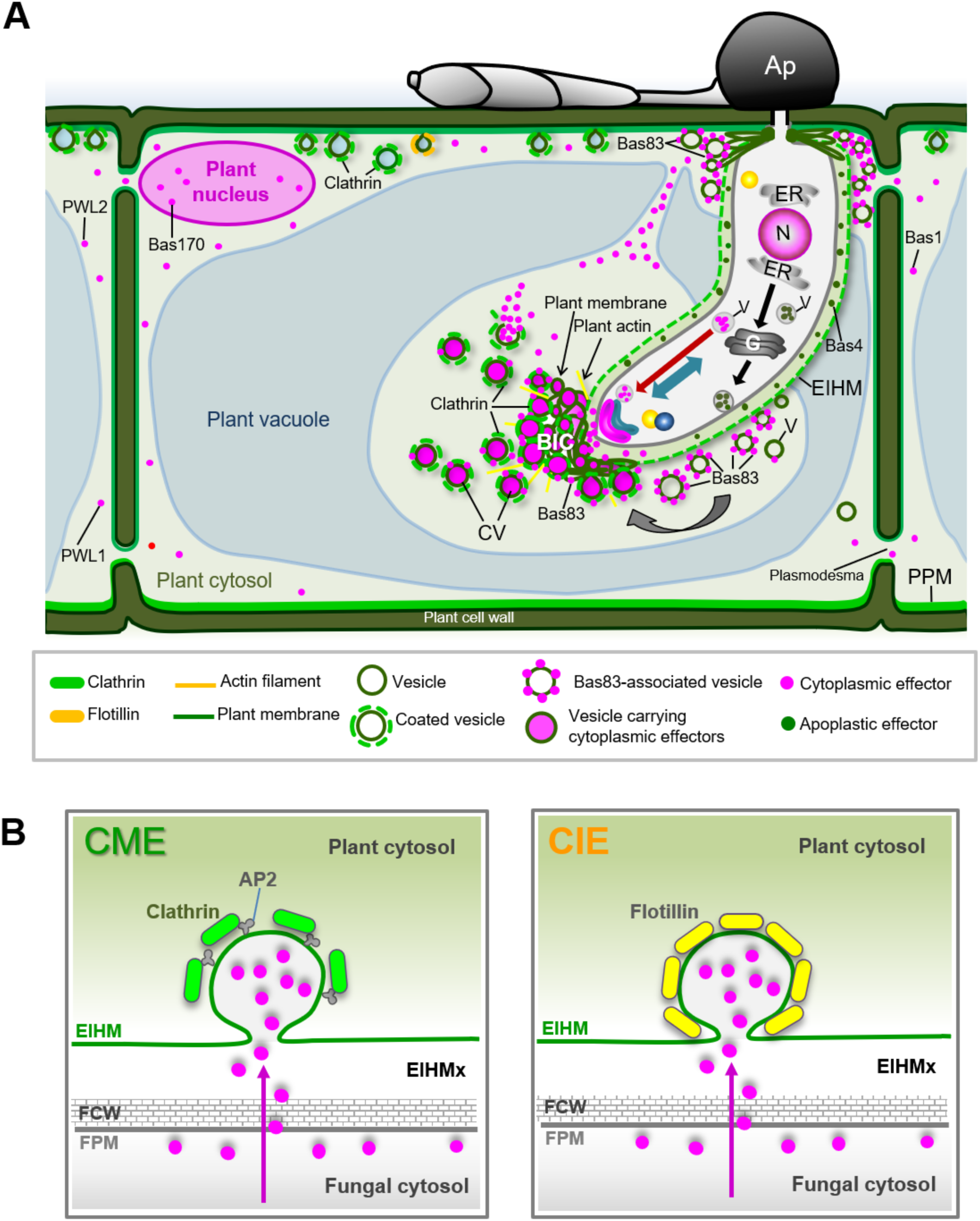
Model for *M oryzae* effector secretion and translocation into rice cells. **A.** This cartoon represents the specialized Biotrophic Interfacial Complex at the growing tip of primary hyphae (tip-BIC) at 22-24 h.p.i. (hours post-inoculation), but similar mechanisms apply to side-BICs left behind after differentiation of the first bulbous invasive hyphae (IH). At successful infection sites, apoplastic effectors (dark green circles), including Biotrophy-associated secreted protein-4 (Bas4), are secreted from the PH by conventional golgi-dependent secretion (black arrows), which is blocked by treatment with Brefeldin A. Apoplastic effectors accumulate in the separate compartment enclosed by the EIHM (inside the dark green dashed line around the PH). In contrast, fluorescently-labeled cytoplasmic effectors (red circles), including Pwl2, Pwl1 and Bas1, are secreted into BICs by a nonconventional, Brefeldin A-insensitive secretion system involving the exocyst and SNARE protein Sso1 (red arrows). Cytoplasmic effectors also accumulate inside the EIHM compartment around the BIC-associated cells, the PH and first differentiated bulbous IH cell (not shown). In contrast, cytoplasmic effectors do not accumulate in the EIHM surrounding nonBIC-associated hyphal cells, which are tightly surrounded by the host vacuole. Our current study expands this model by showing that cytoplasmic effectors are packaged in dynamic vesicles inside BICs and they can sometimes be observed in the host cytoplasm surrounding BICs, and that these effector vesicles co-localize with fluorescent plant plasma membrane markers LTi6B:GFP and FM4-64, as well as clathrin-mediated endocytosis (CME) marker CLC1:eGFP. These co-localization studies, additional localization with plant actin in BICs, and functional analyses via VIGS and pharmacological inhibitors support CME as a major mechanism for effector translocation. Further research should focus on two novel cytoplasmic effectors we describe. Bas83 labels empty membrane vesicles, possibly recruiting more host membrane to the BIC and the EIHM surrounding BIC-associated cells. Bas170 localizes to effector vesicles directly under the appressorium and accumulates in the host nucleus between BICs and the host cell periphery would facilitate movement of translocated effectors to the host nucleus and other sites of action in the host cell, as well as movement of cytoplasmic effectors through plasmodesmata into surrounding host cells to prepare them for invasion. Key: N, fungal nucleus; ER, endoplasmic reticulum; G, Golgi apparatus; V, transport vesicle. **B.** Working models for vesicle formation and effector translocation in the rice-*M. oryzae* interaction through CME or clathrin-independent endocytosis (CIE). Although our current work supports a role for CME, some involvement of CIE in effector translocation cannot be ruled out as playing a minor role. AP2, Adaptor Protein-2 complex; EIHM, extrainvasive hyphal membrane; EIHMx, extrainvasive hyphal matrix; FPM, fungal plasma membrane; FCW, fungal cell wall.

Both VIGS and pharmacological inhibitor strategies have potential interpretation problems. Rice plants that received the CHC-1 and APC-2 silencing vectors were significantly reduced in rice blast symptoms, compared to the empty vector control and to Flot1-silenced lines (Figure 7; Supplemental Figure S7). However, this could be due, at least in part, to inevitable health costs to the plant through silencing component genes for essential cellular processes. Unhealthy rice plants are generally less susceptible to blast disease (Kato, 1974). Additionally, care must be taken in the interpretation of results in pharmacological studies based on new evidence that endocytosis inhibitors characterized in mammalian systems may not have the same targets in plants (Dejonghe et al., 2016; Dejonghe et al., 2019), and pharmacological agents also have off-target effects (Hicks and Raikhel, 2010; Dutta and Donaldson, 2012; Dejonghe and Russinova, 2017). However, the consistent swollen BIC phenotype, seen only with treatments that inhibit CME and consistent with trapping of cytoplasmic effectors inside the BIC (Figures 7-9), lends credibility to our interpretation of these results. Our results do not rule out a minor role for CIE in effector translocation because CIE inhibition has some impact. It remains to be determined if this impact is due to some minor role for CIE or to the reported minor impact of CIE inhibitors filipin and methyl-ß-cyclodextrin on CME (Rodale et al., 1999; Dutta and Donaldson, 2012).

We report that sizes and content of effector vesicles are dynamic through time as host cell invasion progresses. At our levels of resolution, effector vesicles are relatively smaller (median diameter 249 nm) in tip-BICs and increase in size (median diameter 647 nm) especially in later stage side-BICs (Figure 2D). These results confirm and expand on a previous report that cytoplasmic effectors occur in ~500 nm vesicles in the outer BIC domes beside bulbous IH (Mochizuki et al., 2015; Nishimura et al., 2016; Nishizawa et al., 2016). It is unknown why these pathogen-induced effector vesicles have so much larger diameters than reported for plant clathrin coated vesicles (CCVs), which have mean diameters of 59.1 nm for the pentagonal basket population, of 66.1 nm for the hexagonal basket population, and of 85 nm for the irregularly ordered basket population (Narasimhan et al., 2020). The clear increase in effector vesicle sizes through time could be due to biogenesis of larger vesicles, or to aggregation and/or fusion of effector vesicles. Narasimhan et al. (2020) report that Arabidopsis CCVs show delayed and sequential uncoating, and they present evidence for aggregated, partially fused CCVs associated with internal membranes. Future research should explore the differences and similarities in the biology of effector vesicles and normal plant CCVs.

We have identified additional factors that may play a role in effector translocation. We report accumulation of actin in the BICs, as detected in stable transgenic plants expressing LifeAct:eGFP and by staining with rhodamine-labeled phalloidin (Figure 5). It was recently reported that actin is not required for internalization of clathrin-coated vesicles in plants as it is in the well-studied systems in mammals and yeast (Qi et al., 2018; Johnson et al., 2020; Narasimhan et al., 2020). Although a role for actin in plant endocytosis is unknown, actin filaments may be involved in movement of vesicles and cargo sorting after internalization. We hypothesize that actin filaments may be involved in short-distance transport of effector vesicles in the rice cell cytoplasm. Additionally, we identified a novel effector, Bas83, that labels non-effector vesicles and EIHM focused around BICs and the BIC-associated cells. Future research will determine if Bas83 could play a role in replenishing host membranes needed for extensive endocytotic activity in the BIC (Figure 4; Supplemental Figure S4).

We report two new lines of evidence that effectors are being secreted and translocated into the host cytoplasm from appressoria even before obvious growth of primary hyphae inside host cells. First, we identified an effector, Bas170, that naturally accumulates in host nuclei before visible growth of primary hyphae, and it also localizes to vesicles underneath appressoria at this early time point. Second, effector Pwl2:mRFP, which is not normally observed before the tip-BIC stage, accumulates under appressoria that failed to penetrate after both VIGS silencing and chemical inhibition of rice CME. These new findings are consistent with reports that rice cells recognize and respond to *M. oryzae* before and independently of appressorial penetration (Xu et al., 1998). It has been predicted that *M. oryzae* appressorial pores, a transient cell wall-less region of the appressorium adjacent to the plant cuticle, might be involved in molecular communication between pathogen and host before penetration (Howard and Valent, 1996). Previously, the only *M. oryzae* effector that was a candidate for secretion through the appressorial pore was a presumed secondary metabolite that is produced by the *AVR* gene *ACE1* encoding a hybrid polyketide synthase (PKS) and nonribosomal peptide synthetase (NRPS) that is specifically expressed in the cytoplasm of appressoria before and during penetration (Fudal et al., 2007; Collemare et al., 2008). Therefore, our new effector localization results suggest similar exquisite staging of effector expression as reported for the crucifer pathogen *Colletotrichum higginsianum*, which secretes ChEC effectors under the appressorial pores and at subsequent stages during the invasion of *Arabidopsis thaliana* cells (Kleemann et al., 2012). Specifically, this fungus focally secreted effectors Chec6 and Chec36 from the appressorium pore, suggesting these effectors may be translocated before fungal penetration (Kleemann et al., 2012). Our results with Bas170 together with ChECs and Ace1 protein localization patterns indicate that many effectors might be focally secreted through appressorial pores (Fudal et al., 2007; Kleemann et al., 2012), highlighting a potential role for fungal appressoria in effector delivery. Recently, Yan et al. (2022) classified 10 modules of temporally co-expressed genes ranging from 0 hpi to 144 hpi, including some localized to the appressorial pore, among 546 predicted MEP (*Magnaporthe* effector protein) genes.

The effector vesicle dynamics that we report support the idea that front-loaded effector translocation occurs during the early stages of host cell invasion. Specifically, tip-BIC vesicles are relatively small, abundant and selective in terms of sorting different cytoplasmic effectors, Bas1:eYFP and Pwl2:mRFP, into different vesicles. In contrast, side-BICs contain fewer, larger, less-selective vesicles that frequently contain both Bas1:eYFP and Pwl2:mRFP. Effector vesicles observed in the cytoplasm were almost uniformly found at the side-BIC stage, perhaps indicating a slow-down of the mechanism for effector release from the vesicles as BICs mature. Indeed, rare very large vesicles can be seen in some host cells at later stages of invasion (Figure 2C). The mechanism for forming vesicles may result in larger vesicles, or vesicles may fuse before releasing effectors to the cytoplasm in the later BIC stages. Slowdown of the translocation system would appear consistent with previous reports of loss of activity in BICs as host cells fill with IH (Khang et al., 2010; Jones et al., 2021). Front-loading effector translocation from BICs would make sense because cell-to-cell movement of blast cytoplasmic effectors through plasmodesmata must occur in the early stages of host cell invasion while the EIHM enclosing the IH remains intact and plasmodesmata in the invaded cell remain open (Sakulkoo et al., 2018; Jones et al., 2021). Active BICs are surrounded by host cytoplasm with dynamic connections to the cell periphery (Khang et al., 2010; Supplemental Movie S1), where cytoplasmic streaming would rapidly disperse effectors to pit fields containing plasmodesmata. In contrast, nonBIC-associated IH cells invaginate the host vacuole and grow in close proximity to the host vacuolar membrane with little surrounding cytoplasm (Mochizuki et al., 2015; Jones et al., 2021). Specifically, the host vacuole remains intact, but shrinks as IH grow. Both the EIHM and vacuolar membrane become disrupted, indicating death of the invaded cell, around the time IH move into neighboring cells to repeat their BIC-mediated invasion strategy (Mochizuki et al., 2015; Jones et al., 2021).

There is evidence supporting occurrence of clathrin-mediated endocytosis at the EHM in one additional pathosystem. That is, transmission electron micrographs of the interfacial matrix between the intracellular hyphae [called monokaryotic (M)-haustoria] of the monokaryotic rust fungus *Uromyces vignae* and its host cowpea (*Vigna unguiculata*) shows clathrin-coated pits (diameter ca. 50-70nm) on the EHM at M-haustoria hyphal tips that contained dense cytoplasm and were presumably growing (Stark-Urnau and Mendgen, 1995). The coated vesicles were labeled by an antibody recognizing the clathrin heavy chain subunit. Long tubules extending into the host cytoplasm, as well as coated vesicles in the surrounding host cytoplasm, were also labeled with the antibody, indicating that endocytosis occurs at the EHM (Stark-Urnau and Mendgen, 1995; O’Connell and Panstruga, 2006). It was suggested that these endocytic vesicles might be involved in effector uptake into host cells in addition to membrane recycling from host exocytosis. Unfortunately, lack of transformation capability and identification of effectors makes it difficult to follow up on these intriguing results with *U. vignae*. Additional TEM studies support our results that transient silencing of lipid raft-mediated endocytosis component Flotilin1, as well as filipin treatment, had little effect on BIC structure and effector vesicle formation. Freeze-fracture transmission electron microscopy (TEM) following filipin treatment revealed an absence of granular filipin-sterol complexes on the EHM of two rust fungi, *Puccinia coronata* and *Uromyces appendiculatus* (Harder and Mendgen, 1982), suggesting that the extrahaustorial membrane contains less sterol than normal plasma membranes, and indicating that the EHM appears depleted in sterol-rich lipid rafts.

Differences in infection strategies among various filamentous biotrophic or hemibiotrophic pathogens would be consistent with use of different strategies for effector translocation in different pathosystems. A recent report indicates that a complex of seven proteins from the smut pathogen *U. maydis* is critical for infection and implicated in translocation of cytoplasmic effectors into maize cells (Ludwig et al., 2021). This basidiomycete fungus colonizes host tissue as intracellular hyphae that pass through and between host cells without obviously specialized interfacial zones, and fluorescent fusion proteins of cytoplasmic effectors have not been visualized in the host cell cytoplasm in this pathosystem (Lo Presti et al., 2017). *U. maydis* produces galls on maize seedlings and grows systemically as the plant grows to finally invade ovaries, which swell into dramatic sporulating galls. In contrast, *M. oryzae* densely colonizes a localized tissue area that will become a visible eyespot lesion that releases spores to reinitiate the infection cycle within 7 days. Different pathogenic lifestyles could present different demands for cytoplasmic effector translocation.

Taken together, our data strongly suggest that clathrin-mediated endocytosis occurring in BICs is a key mechanism for internalization of *M. oryzae* effectors inside living rice cells, beginning even before appressorium penetration of the host surface. Further confirmation will come through continuing research to determine how the endocytic machinery is recruited to BICs and how it interacts with effector cargos (Qi et al., 2018). It is important to understand if, and if so, how, Bas83 contributes to membrane recruitment to BICs and the EIHM surrounding BIC-associated cells. It is of interest to understand how supposedly coordinately-expressed effectors Bas1:eYFP and Pwl2:mRFP can be sorted into different effector vesicles, as seen in Figure 2A. It is important to understand vesicle dynamics, including and why effector vesicles become larger and more persistent as BICs mature, and why effector vesicles are more easily observed in the host cytoplasm near-by maturing side-BICs (Figure 2). Identification of effectors with critical roles in the translocation process is a high priority, including effectors that are involved in disrupting effector vesicles and releasing effectors into the rice cytoplasm. It is important to understand the role for appressoria in effector secretion before penetration, and how these early effectors, such as the host nuclear localized effector Bas170, function to promote infection. Finally, it is critical to translate molecular mechanisms of blast biotrophic invasion into strategies for controlling devastating diseases on rice, wheat and other food crops worldwide.

## METHODS

### Live-cell imaging of *M. oryzae* effectors *in planta*

Fungal strains were stored in dried filter papers at −20°C, and cultured on rice polish agar plates at 25°C under continuous light for 2 weeks (Valent and Chumley, 1991). Rice sheath inoculations were performed as described (Kankanala et al., 2007) with the following modification. We used sheath pieces that were thickly trimmed (~7 rice cell layers thick) compared to thinner trimmed sheaths (~3 cell layers thick) in previous publications. Brightfield images were less detailed, but the endocytic machinery appeared more active, providing enhanced microscopic resolution of fluorescent marker dynamics. Susceptible rice variety YT-16 was used unless mentioned otherwise. Briefly, 7-cm-long sheath pieces from 3-week-old plants were placed in a sealable Pyrex glass moist chamber. Leaf sheath sections were placed on inverted 8-well PCR tube strips to avoid contact with wet paper and to hold epidermal cells directly above the mid-vein horizontally flat for uniform inoculum distribution in the trimmed sheath pieces (Oliveira and Valent, 2021). A spore suspension (10^4^ spores/ml in sterile 0.25% gelatin, Cat. #G-6650, Sigma-Aldrich) was prepared from 10-day old cultures and was injected into one end of the sheath using a 100-ml pipette. Each segment was trimmed at 18–30 h.p.i., treated with specific dyes or inhibitors, or imaged immediately by laser confocal microscopy. Biological replicates were independent experiments performed with fungal cultures fresh out of frozen storage and with new rice plants. All conclusions are supported by at least 3 biological reps, with each replication including observation of around 100 infection sites.

Confocal imaging was performed with a Zeiss LSM780 confocal microscope system using two water immersion objectives, C-Apochromat 40x/1.2 WCorr. And C-Apochromat 63x/1.2WCorr. Excitation/emission wavelengths were 488 nm/505–550 nm for eGFP and FM1-43, and 543 nm/560–615 nm for mRFP, mCherry, Phalloidin and FM4-64. Image acquisition and processing, some vesicle quantification, and fluorescence intensity linescans were done using Zeiss ZEN 2010 software. Images of effector vesicles in Figure 3C were obtained using a Leica SP8 confocal microscope system with a water immersion objective HC PL APO 63x/1.20 WCorr. Excitation/emission wavelengths were 488 nm/505–550 nm for eGFP and 543 nm/560–615 nm for mRFP. Leica SP8 software at the University of Exeter and at the Lousiana State University Agricultural Center were used for some vesicle quantification and for analyzing ES9 and ES9-17 CME inhibitor assays.

### Size measurements of vesicles carrying effector and clathrin fluorescence

To measure BIC vesicle sizes, high resolution confocal microscopy images of BICs were generated from infected YT16 rice undergoing infection of *M. oryzae* strains KV170 or KV209 expressing Bas1:mRFP or Pwl2:mRFP, respectively. We began with maximum intensity projections of Z-stack image series of BICs obtained with the Zeiss and Leica confocal microscopes. However, when vesicles were too close in the collapsed images, distinction between two or more vesicles was attempted using individual images in the Z-stack. For example, see Supplemental Movie S3. Scale bars were set to 1 µm. Amplified images were opened in ImageJ (Schneider et.al., 2012) and distances in pixels was correlated to the 1 µm (1000nm) scale bars. Vesicle diameters were measured using the length measurement tool. Data was exported in Excel and analyzed using the Prism 9 software.

### Fungal strains, DNA manipulation, and fungal transformation

*Magnaporthe oryzae* wild type strain Guy11, a field isolate from rice in French Guiana, was obtained from J.L. Notteghem (Centre de Cooperation Internationale en Recherche Agronomique pour le Developpment, France). *M. oryzae* transformants are described in Supplemental Table S1 online. Effector:mRFP and Effector:eGFP expression plasmids were constructed by PCR amplifying different effector gene regions and fusing them to the N-terminus of mRFP and eGFP, respectively. Details of construction of each plasmid are described in Supplemental Table S2. All primers used are listed in Supplemental Table S3. For all fusion constructs, transcriptional and translational fusions were verified by DNA sequencing. Plasmids were transformed into laboratory strain Guy11 (Leung et al., 1988) using *Agrobacterium tumefaciens*–mediated transformation (Khang et al., 2006). In several cases, two fluorescently labelled effectors were introduced by co-transformation with separate plasmids (Sweigard et al., 1995). For each plasmid construct, 10 independent transformants were assayed for fluorescence intensity during host invasion and for consistent localization patterns. Transformants with high fluorescence intensities were stored and studied further (see Supplemental Table S1 online). For each construct cloned in *M. oryzae*, at least two independent transformants showing identical sporulation, growth and infection phenotypes to the wild type were selected and assayed together.

### Strategies for targeted deletion of Bas83 gene

*Split-marker recombination method for Bas83 gene knockout via protoplast transformation*: Targeted gene replacement mutation of the *M. oryzae Bas83* gene was attempted using the split marker strategy as modified by Kershaw and Talbot (2009). Gene replacement was performed by replacing the 606-bp *Bas83* locus with a hygromycin resistance selectable marker, encoding a 1.4-kb hygromycin phosphotransferase (*HPH*) resistance cassette (Carroll et al., 1994). The two overlapping segments of the *HPH* templates were PCR amplified using primers M13F with HY and M13R with YG (Catlett et al., 2003) (see Supplemental Table S3 online) as described previously (Kershaw and Talbot, 2009). One-kb DNA fragments upstream and downstream of the *Bas83* open reading frame were generated using the primers Bas83LF-F and Bas83LFHY-R and Bas83RFYG-F and Bas83RF-R amplified from genomic DNA of the Guy11 strain. A second-round PCR reaction was performed to fuse the overlapping split hph marker templates with the left and right flanking regions of the Bas83 locus. Wild-type *M. oryzae* strain Guy11 was then transformed with the deletion cassettes (2 μg of each flank). Putative transformants were selected in the presence of hygromycin B (200 μg/ml) and analyzed by PCR using the primers Bas83:BASKOtest-F and Bas83:BASKOtest-R (Supplemental Table S3; Supplemental Figure S5). This assay was repeated three times. Gene sequences on either side of *Bas83* were retrieved from the NCBI database (https://www.ncbi.nlm.nih.gov/).

*Attempted Bas83 deletion via Agrobacterium-mediated transformation*: For gene replacement transformation, cassettes were constructed by amplifying ~1.0 kb of 5′- and 3′-flanking regions for each predicted coding sequence. The *HPH* gene (Carroll et al., 1994) was cloned between the two flanking regions using a fusion PCR strategy. The three pieces together were cloned first into the pJET1.6 vector (Invitrogen) for sequence analysis and later into binary vector pGKO2 (Addgene: Plasmid #63617; Khang et al., 2005) using a restriction ligation strategy. *M. oryzae* KU70 and KU80 spores were transformed using *A. tumefaciens* (Khang et al., 2005) strains AGL1 and EHA105 (Hellens et al., 2000). After two rounds of selection in TB3 media containing 200 μg/mL of hygromycin, 400 independent fungal transformants were analyzed for gene replacement events by PCR amplification using the primers Bas83:BASKOtest-F and Bas83:BASKOtest-R (Supplemental Figure S4). This assay was repeated three times.

### Staining with FM4-64, FM1-43, and Rhodamine Phalloidin, and treatment with pharmacological inhibitors

To examine membrane dynamics at the BIC, rice sheaths (cv. YT16) were inoculated with *M. oryzae* transformants expressing fluorescently-labeled effectors (2×10^4^ spores/ml in 0.25% gelatin solution) and incubated for 20-26 hours at 25°C for subsequent FM-dye treatment. Infected rice sheaths were treated with FM4-64 or FM1-43 (4 mg/ml in water). An aqueous 17mM stock solution of FM4-64 (Cat #13320, Invitrogen, Carlsbad, CA) and FM1-43 (Cat # T3163, Invitrogen, Carlsbad, CA) were prepared and stored at −20°C as described (45). *In planta*, inoculated leaf sheaths, 24 h.p.i., were hand-trimmed into thin sections (3-7 layers of cells; 10 x 10 mm) and incubated in a 10-mM aqueous working solution for 3–5 hours. Leaf sheaf sections were subsequently transferred to ultrapure water and incubated for 20 min (25 °C) to remove excess dyes for uniform membrane staining. For *in planta* experiments with Rhodamine Phalloidin (Cat# R415, ThermoFisher Scientific), we prepared working solutions of 66 µM in 0.1% DMSO and treated the inoculated tissue as described for FM dyes.

To examine the effects of endocytosis inhibitors on *in planta* effector uptake, inoculated trimmed rice sheaths were incubated at 25°C with Endosidin9 (ES9) (10µM), or Endosidin9-17 (ES9-17) (30µM), or Methyl-β-cyclodextrin (20 mM), or Chlorpromazine (10 µg/mL), or Cantharidin (25 mM), or Fluazinam (20 mM), or Triclosan (20mM), or Wortmannin (20 nM), or Concanamycin A (50 mM) (all from Sigma) solution for 1-5 hours. Negative controls were performed with ultrapure water (Methyl-β-cyclodextrin, Chlorpromazine, and Fluazinam) or 0.1% DMSO (ES9, ES9-17, Cantharidin, Filipin and Wortamanin) (see Table 1).

### Generation of transgenic rice plants

Production of transgenic plants expressing the rice plant plasma membrane marker low-temperature inducible protein 6B, LTi6B:GFP was previously described (Mentlak et al., 2012; Giraldo et al., 2013). Transformants of rice cultivar Sasanishiki expressing LifeAct:eGFP were generated in the laboratory of NJT. Expression plasmids for CME and CIE marker genes were constructed by PCR amplifying the marker gene regions and fusing them to the N-terminus of eGFP. The *CLC1* gene was amplified from genomic DNA of YT16 rice using the primers KpnI_OsCLC1Prom1-F1 and XhoI_OsCLC1-R1 (See Supplemental Table S3). The PCR product was digested with *Kpn*I and *Xho*I and integrated into pSH1.6EGFP (Plasmid #42323, Addgene). The *Flot1* gene was amplified from genomic DNA from YT16 rice using the primers HindIII_OsFlot1Prom1-F1 and Eco47III_OsFlot-R1. The PCR product was digested with *Hind*III and *Eco*47III and integrated into pSH1.6EGFP. CLC1:eGFP was amplified from pSH1.6_CLC1:eGFP using the primers CACC_OsCLC1-F1 and eGFPT2-R2, and cloned into pENTR (pENTR™/D-TOPO™ Cloning Kit, ThermoFisher Scientific) following the manufacturer’s protocol. Flot1:eGFP was amplified form pSH1.6_Flot1:eGFP using the primers CACC_OsFlot1-F1 and eGFPT2-R2, and cloned into pENTR following manufacturer’s protocols. The endocytosis marker:eGFP constructs were cloned into the *Agrobacterium* vector pIPKb001 (Himmelbach et al., 2007) using the Gateway LR Clonase II kit (Invitrogen) following the manufacturer’s protocol. All the fusion constructs had transcriptional and translational fusions verified by DNA sequencing. Gene fusions for expressing the rice endocytosis markers CLC1:eGFP (CME) and Flot1:eGFP (CIE) under their native promoters were introduced into *Agrobacterium tumefaciens* EHA105 using the freeze-thaw method (Holsters et al., 1978). Mature seed-derived callus from rice (*Oryza sativa L*. *Japonica*) cv. YT16 was used for *Agrobacterium*-mediated transformation (Park et al., 2001). After inoculating with *A. tumefaciens*, callus was transferred to regeneration medium for 4-10 weeks at 25°C under a 16-h photoperiod. The regenerated shoots were transferred to rooting medium for four more weeks, then established in soil. Putative transformants of rice were selected on 100 µg/ml hygromycin, and expression checked by qRT–PCR and epifluorescence microscopy. Thirty independent T1 transgenic lines with CLC1:eGFP and 28 independent T1 lines with Flot1:eGFP were recovered. All lines were assayed by microscopy for fluorescence pattern and for fluorescence intensity. All lines for each construct showed the predicted fluorescence pattern. We chose lines with the highest levels of fluorescence for assays.

### Silencing of rice genes

In order to evaluate the role of plant endocytosis on disease development and effector uptake, we performed VIGS using the Brome mosaic virus system developed by R.S. Nelson (Noble Foundation, Ardmore, OK). We targeted *AP-2α*, *CHC1* (CME) and *Flot1* (CIE) mRNA in rice plants (cultivar IR64) undergoing infection. The target rice sequences were cloned in the pC13/F3-13m VIGS vector as described (Ding et al., 2006; Ding et al., 2007) and subsequently sequenced to verify sequence accuracy. RNA transcripts were synthesized from the final VIGS vector. An empty vector pC13/F3-13m was used as negative control. The final infectious transcripts of pC13/F3-13m / AP-2α, pC13/F3-13m / CHC1, pC13/F3-13m /Flot1 and pC13/F3-13m (a control virus without an insert) were inoculated with pC13/F1+2 in 7 to 10 days old IR64 rice plants according to protocols described by Ding et al (2006). Knockdown of the target genes was confirmed by qRT-PCR at 15 days post inoculation. At this point, rice plants showing satisfactory reduction of the transcription levels were used in our standard conidial spray inoculation and leaf sheath assays with fungal strains expressing fluorescently-labeled effectors.

### Statistics

All experiments were performed with at least 3 biological replications, which are independent experiments with fungal cultures directly growing out from frozen storage and different rice plantings. Biological replications included at least two technical repeats (independent assays with the same biological materials) for further confirming reproducibility of the data. In all cases, technical and biological replications gave consistent results. Sample sizes, number of biological replicates, and the statistical tests used in each experiment are specified in the figure legends. Data were analyzed using an unpaired two-tailed Student’s t-test. P=0.05 was considered non-significant and exact values are shown where appropriate. All statistical analysis was performed using R Statistical Software (version 4.1.2) and Prism9 (GraphPad). Dot plots were routinely used to show individual data points and generated using Prism9 (GraphPad). Bar graphs show the mean±s.e.m. (unless stated otherwise) and were generated using Prism9 (GraphPad). Analysis of non-normal datasets are represented by box-and-whisker plots that show the 25th and 75th percentiles, the median indicated by a horizontal line, and the minimum and maximum values indicated by the ends of the whiskers.

### Accession Numbers

Sequence data for genes of *M. oryzae* used in this article can be found in the GenBank/EMBL database under the following accession numbers: BAS1, FJ807764.1/NC_017844; PWL1, U36923.1; PWL2, U26313.1/NC_017853; Bas4, FJ807767.1/NC_017852.1; BAS83, MGG_08506.6/NC_017851.1; BAS170, MGG_07348.6/NC_017850.1. Rice genes used are: Clathrin Light Chain-1 (CLC1), LOC4337419; Flotilin 1 (Flot1), LOC4348926; Adaptor protein complex-2α (AP-2α), LOC4331370; and Clathrin Heavy Chain-1 (CHC1 gene; LOC4349546.

## ACKNOWLEDGEMENTS

We acknowledge and thank Rick Nelson (Noble Foundation, Ardmore, OK) for providing the VIGS vectors and hosting E.O.-G. in his laboratory to learn the VIGS assay. We thank the Talbot laboratory for assistance and hospitality during a working visit for E.O.-G at the University of Exeter. We thank Hiromasa Saitoh (Iwate Biotechnology Research Center, Kitakami, Iwate, Japan) and Ryohei Terauchi (Iwate Biotechnology Research Center, Kitakami, Iwate, and Kyoto University, Japan) for sharing transgenic rice expressing plant plasma membrane marker LTi6B:GFP. This project was supported by Agriculture and Food Research Initiative Competitive Grant no. #2017-67013-26525 from the USDA National Institute of Food and Agriculture Plant Biotic Interactions program. Additional support came from Louisiana State University Agricultural Center Hatch project #LAB94477 and from Louisiana Board of Regents grant #LEQSF(2022-24)-RD-A-01. This is Contribution no. 22-053-J from the Kansas Agricultural Experiment Station.

## AUTHOR CONTRIBUTIONS

E.O.-G. and B.V. designed experiments and analyzed data: E.O.-G. performed most of the research. T.M.T., J.P. and S.P. produced and analyzed transgenic rice lines expressing the rice CLC1:eGFP and Flot1:eGFP fluorescent fusion protein genes. M.D., M.M.-U., C.R.H. and A.H.V. assisted with microscopy and Bas83 knockout experiments and general laboratory experiments; N.J.T. provided transgenic rice lines expressing LTi6B:GFP and LifeACT:GFP; E.O.-G., B.V. and N.J.T wrote and edited the article.

