## Supplemental Figures and Tables for "Clathrin-mediated Endocytosis Facilitates Internalization of *Magnaporthe oryzae* Effectors into Rice Cells"

SUPPLEMENTAL INFORMATION CONTAINED IN THIS FILE

**Supplemental Figure S1**. BIC effector vesicles are observed with additional fluorescent cytoplasmic effectors. Supports Figure 1A-C.

**Supplemental Figure S2.** Effector vesicles and host translocation at 24 hpi, after hyphal differentiation. Supports Figure 1B.

**Supplemental Figure S3.** Sequence similarities between Bas83 and Bas170. Supports Figure 1D-F.

**Supplemental Figure S4.** Bas83:mRFP localization during invasion of rice cells. Supports Figure 4.

**Supplemental Figure S5.** Strategy for targeted deletion of *BAS83*. Supports Figure 4.

**Supplemental Figure S6.** Rice clathrin light chain-1 uniformly occurs in BICs and can be seen localizing to vesicles labeled by Bas1:mRFP or Bas170:mRFP. Supports Figure 6.

**Supplemental Figure S7.** VIGS silencing of endocytosis components in IR64 rice using the Brome Mosaic Virus system. Supports Figure 7.

**Supplemental Table S1.** *M. oryzae* transformants used in this study.

**Supplemental Table S2.** Plasmids used in this study.

**Supplemental Table S3.** Oligonucleotides used in this study.

**Legend for Supplemental Movie S1.** Optical sections showing a cytoplasmic connection between a BIC and peripheral rice cytoplasm as well as effector vesicles in different cell layers. Supports Figure 1.

**Legend for Supplemental Movie S2.** Optical sections showing effector vesicles in the host cytoplasm and at a distance from a BIC, in tissue stained by endocytosis tracker dye FM4-64. Supports Figure 3D,E.

**Legend for Supplemental Movie S3.** Time-lapse images showing co-localization of clathrin light chain-1 labeled with eGFP and cytoplasmic effector Pwl2 labeled with mRFP in BIC vesicles. Supports Figure 6B.

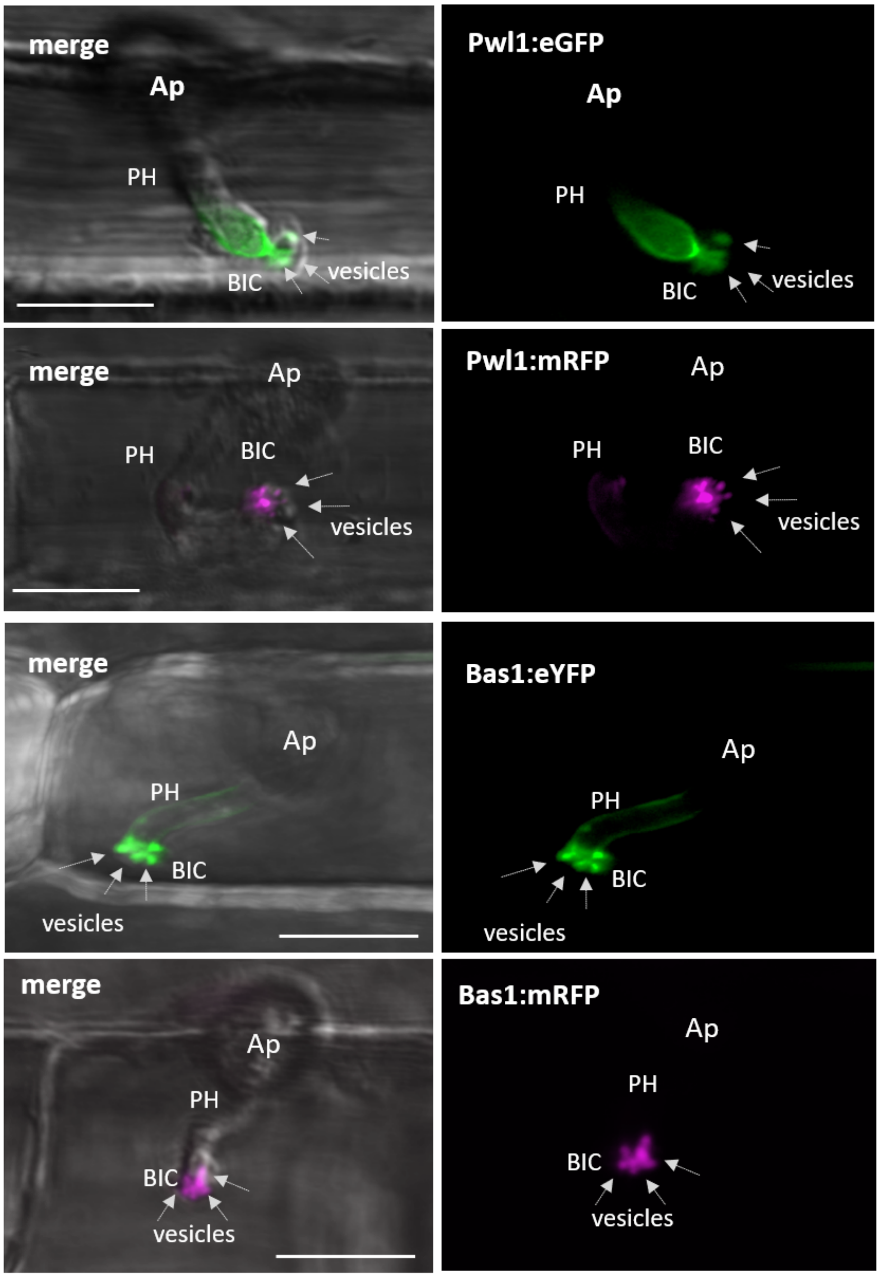


**Supplemental Figure S1. BIC effector vesicles are observed with additional fluorescent cytoplasmic effectors.** Effector vesicles (upper to lower panels) are shown in tip BICs formed by strains KV174 and KV244 expressing Pwl1:eGFP and Pwl1:mRFP, respectively, and by strains KV182 and KV170 expressing Bas1:eYFP and Bas1:mRFP, respectively. These results support Figure 1 obtained with strain KV217 expressing Pwl2:mRFP and Bas4:eGFP, and indicate that effector vesicle formation is independent of cytoplasmic effector and fluorescent protein used. BICs are imaged at the primary hyphal stage at ~19 hpi. Images are merged bright field and eYFP, eGFP or mRFP fluorescence (left) and fluorescence channels alone (right). Bars = 10 µm.

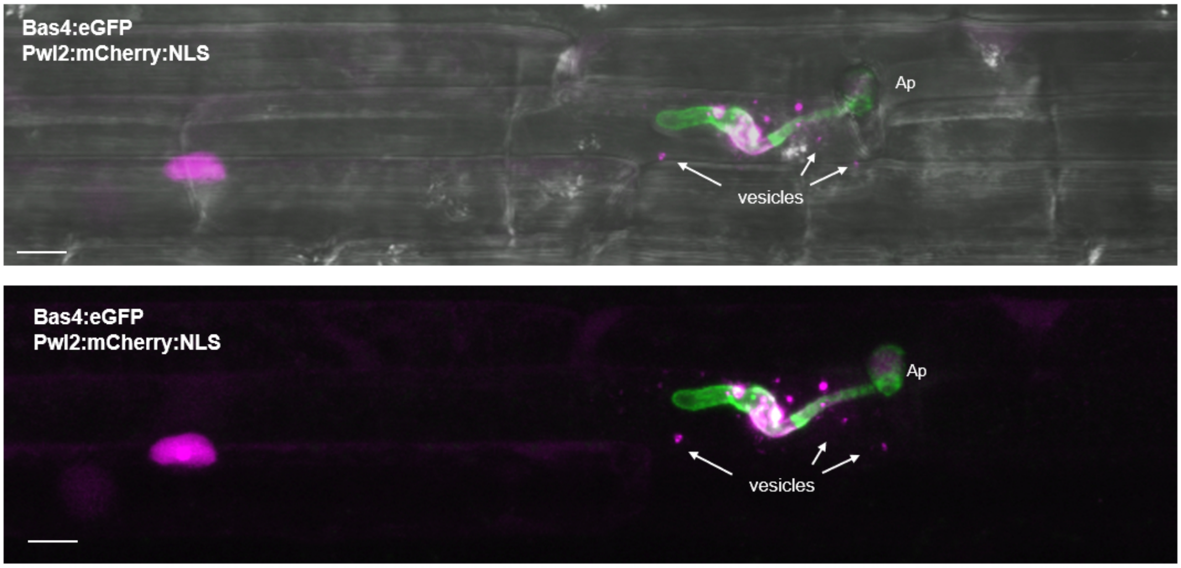


**Supplemental Figure S2. Effector vesicles and host translocation at 24 hpi after hyphal differentiation.** Supports Figure 1B by showing the entire rice cell containing the side-BIC. The BIC, produced by strain KV168 expressing Pwl2:mCherry:NLS (magenta) and Bas4:eGFP (green) at 24 hpi, occurs above the plane of focus on the first bulbous IH cell. Cytoplasmic effector vesicles (white arrows) are visible in the host cytoplasm at a distance from the BIC, and mRFP fluorescence in the host nucleus confirms translocation. The Bas4:eGFP control shows that the EIHM still surrounds the IH. Addition of an artificial NLS to the fluorescent Pwl2 fusion protein facilitates visualization of effector translocation through accumulation in the host nucleus. Images (top to bottom) are merged bright field, eGFP and mRFP fluorescence, and merged eGFP and mRFP. Bars = 10 µm.

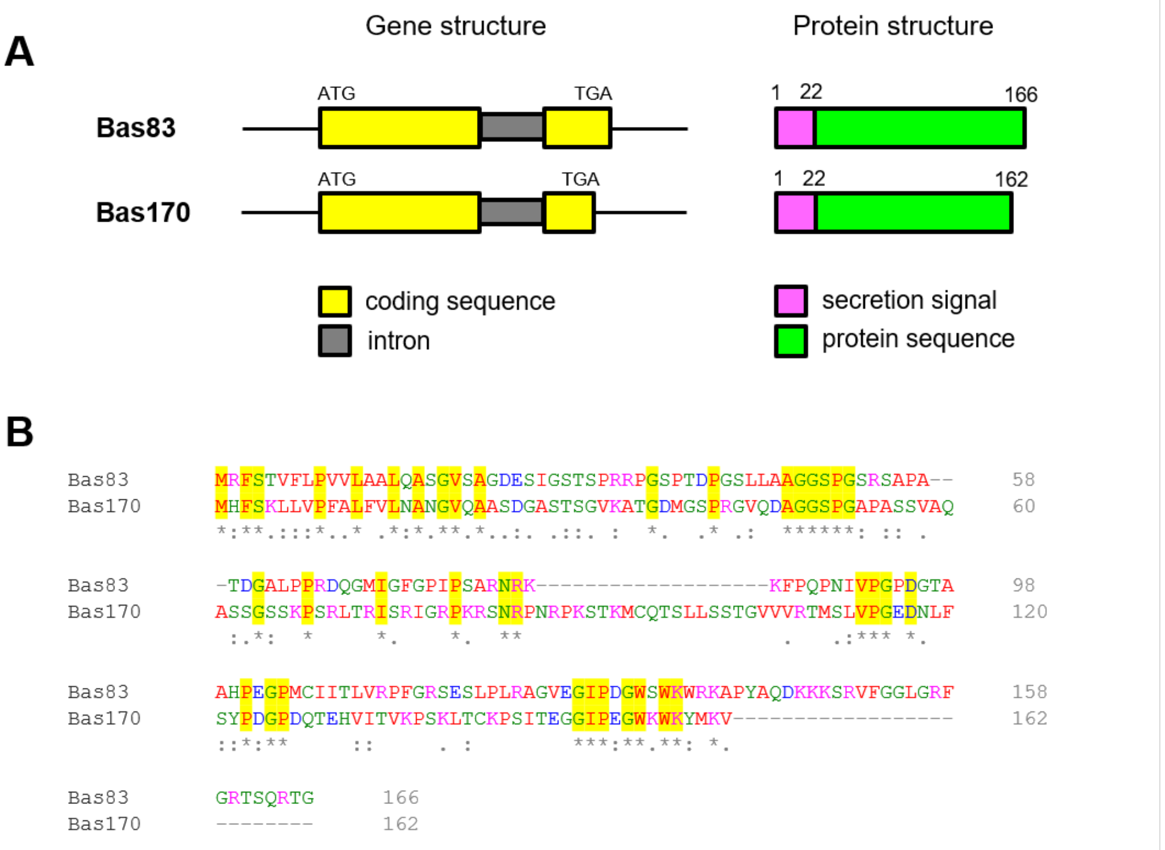


**Supplemental Figure S3. Bas83 and Bas170 show conserved intron positions in their gene structures, but they show low similarity in their protein structure. A.** Gene (left) and protein structure (right) of Bas83 and Bas170 in *M. oryzae* strain 70-15. Exons are shown as yellow bars, introns are grey bars. The predicted proteins show a clear secretion signal (magenta), but no conserved domains (green bars). **B.** Pairwise amino acid sequence alignment between Bas83 and Bas170 proteins shows low levels of sequence similarity.

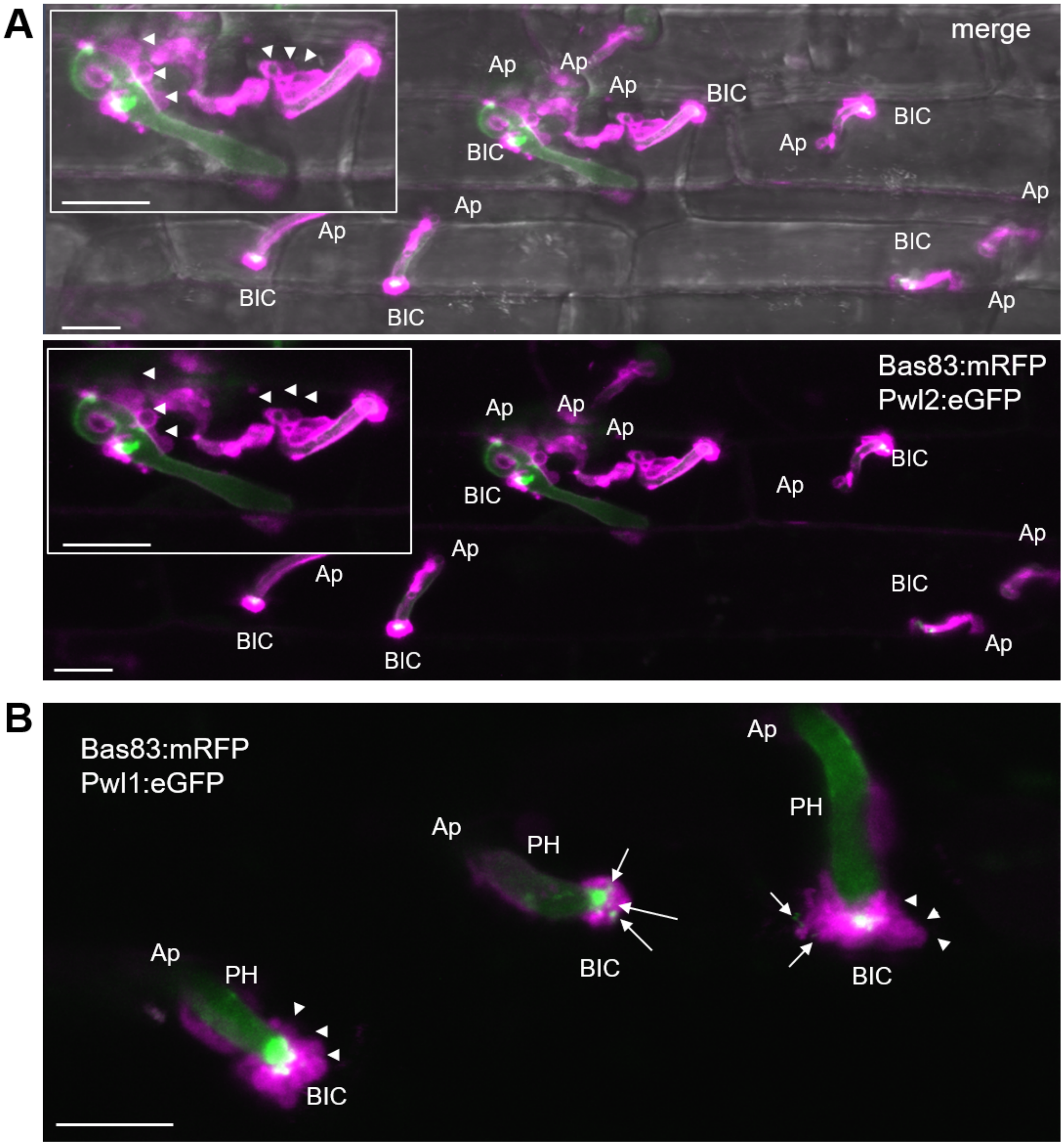


**Supplemental Figure S4. Bas83:mRFP localization during invasion of rice cells.** Multiple infection sites of strains KV222 and KV246 expressing respectively Bas83:mRFP with Pwl2:eGFP or Bas83:mRFP with Pwl1:eGFP in YT16 rice at 22 hpi, complementing data in Figure 4. **A**. Localization of Bas83:mRFP suggests manipulation of plasma membrane near appressoria, primary hyphae and BICs. Insert shows large vesicles (arrowheads) labeled by Bas83:mRFP surrounding biotrophic invasive hyphae. **B.** Localization of Bas83:mRFP in vesicles that lack Pwl1:eGFP fluorescence around the BIC (Arrowheads). Arrows indicate vesicles containing Pwl1:eGFP, perhaps being released from the BIC to rice cell cytoplasm. Images (merged bright field, eGFP and mRFP fluorescence) are shown as projections of confocal optical sections. Bars = 10 µm.

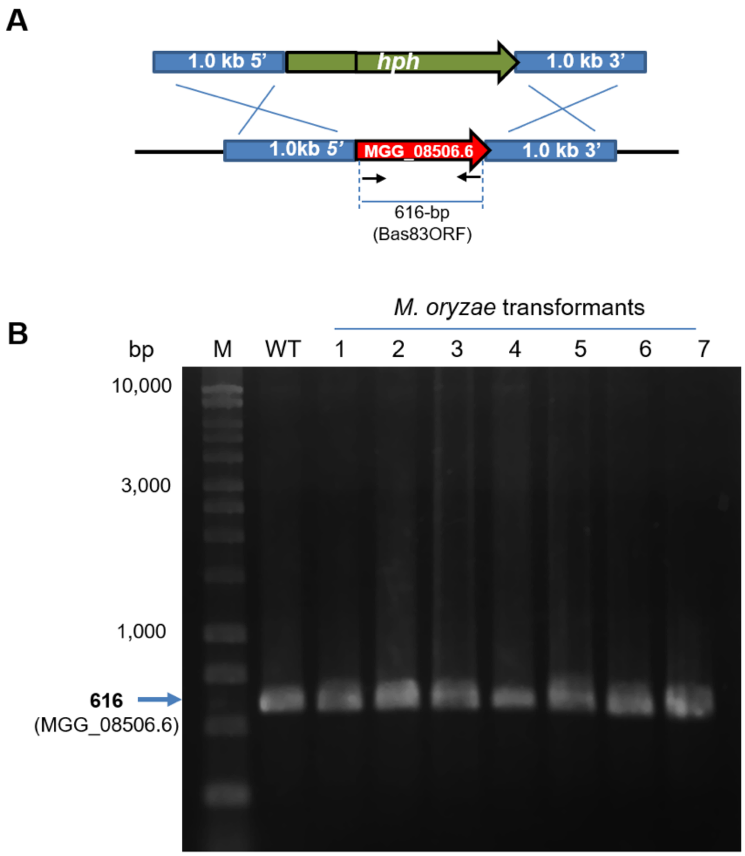


**Supplemental Figure S5. Strategy for targeted deletion of the *BAS83* gene.** Both protoplast-based transformation and *Agrobacterium*-based transformation performed in two laboratories failed to generate knock-out mutations in *BAS83*. **A.** The deletion cassette consisted of the hygromycin cassette cloned between ~1.0 kb 5'- and 3'-flanking sequences of *BAS83*. Arrows indicate binding sites of primers Bas83:BASKOtest-F1 and Bas83:BASKOtest-R1; the bar indicates the predicted ~616 bp-PCR band in the wild-type (WT) strain and transformants with an ectopically integrated deletion cassette. Not to scale. **B.** PCR screen for homologous integration of the deletion cassette. PCR primers (Bas83:BASKOtest-F1 and Bas83:BASKOtest-R1) have binding sites in the coding sequence of *Bas83*, so that the 616 bp band can only be amplified from the wild-type (WT) strain and strains harboring an ectopically integrated deletion cassette. Seven independent transformants (1 – 7) of a total of 400 are shown. PCR analyses showed a ~616 bp band for all transformants tested, indicating ectopic integration of the deletion cassette. M, DNA size marker.

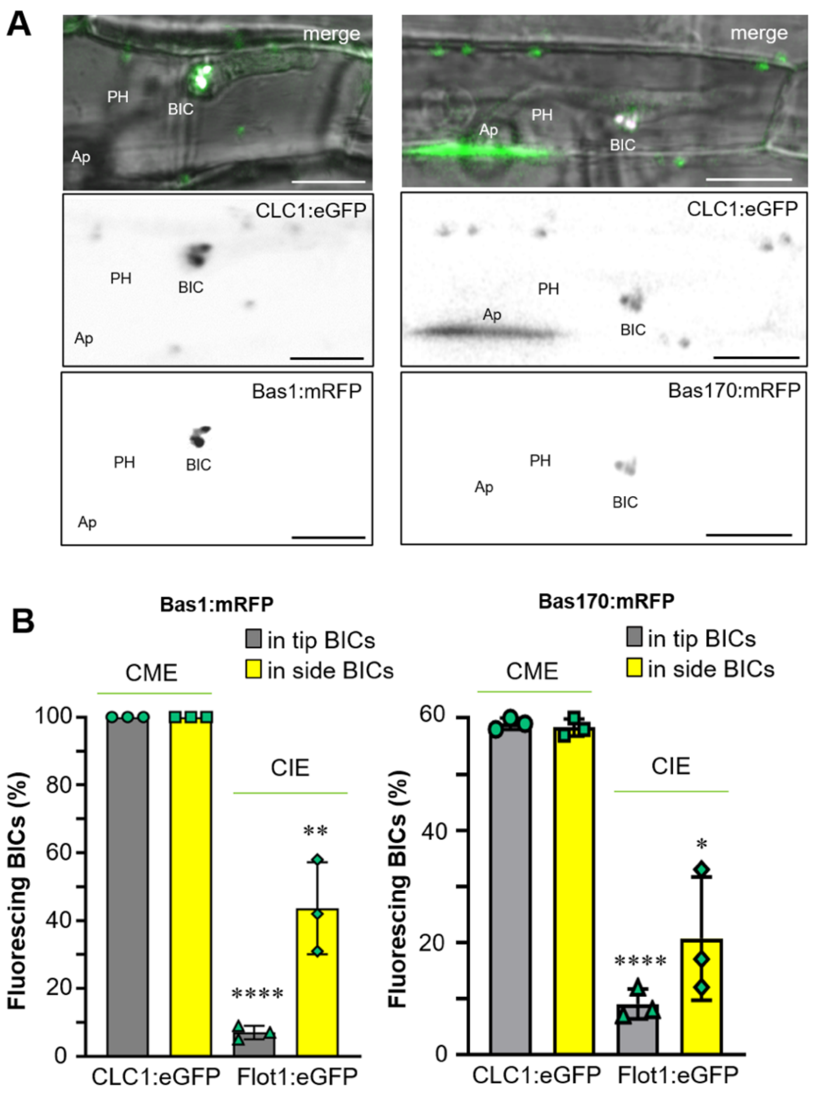


**Supplemental Figure S6. Rice CCLC1:eGFP localizes with BIC vesicles labeled with additional cytoplasmic effectors Bas1:mRFP and Bas170:mRFP.** Supports Figure 6. **A.** Rice leaf sheaths expressing OsCLC1:eGFP were inoculated with strain KV170 expressing Bas1:mRFP (left panels) or strain KV224 expressing Bas170:mRFP (right panels). These infected rice cells contain bulbous IH with late stage BICs. Cell wall autofluorescence is seen below the appressorium in the image with Bas170:mRFP. Images shown top to bottom are merged bright field, eGFP and mRFP (magenta); then eGFP alone and mRFP alone as black and white inverse images. Bars=10µm. **B.** Quantification of colocalization of OsCLC1:eGFP or OsFlot1:eGFP with effector vesicles in tip- or side-BICs. Bars are standard deviations. ****P<0.0001, **P=0.002, *P=0.038; Bas1mRFP/CLC1:eGFP colocalization: three times 98 BICs observed; Bas170mRFP/CLC1:eGFP colocalization: three times 60 BICs observed.

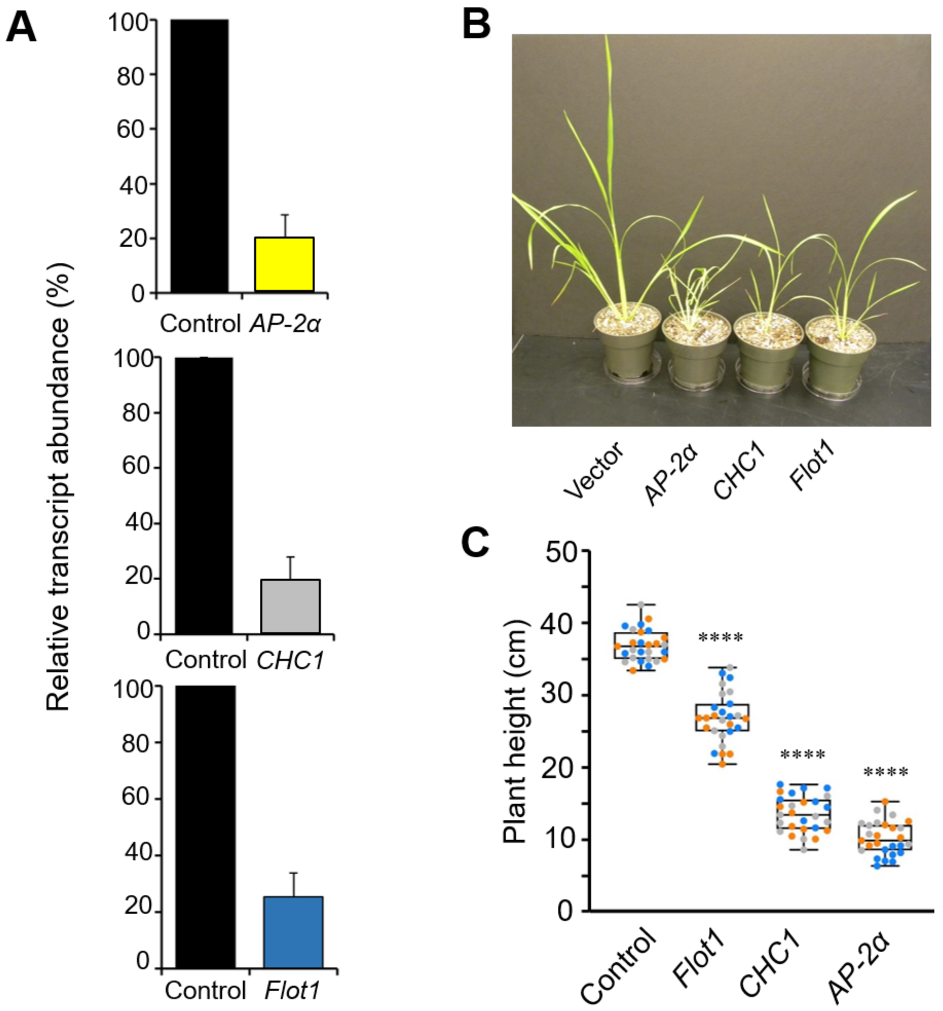


**Supplemental Figure S7. VIGS silencing of endocytosis components in IR64 rice using the Brome Mosaic Virus system. A.** Relative transcript abundance of rice *AP-2*$\alpha$, *CHC1*, and *Flot1* in lines expressing RNAi*AP-2*$\alpha$, RNAi*CHC1*, RNAi*Flot1* silencing constructs relative to vector control plants at 15 days post inoculation. Individual plants showing high levels of silencing were chosen for infection assays. Three biological replications with 9 plants each were evaluated. Error bars indicate standard deviations. **B.** *AP-2*$\alpha$, *CHC1* and *Flot1* silenced plants show significant stunting compared to plants that received the vector control, as expected for inhibiting critical gene functions. **C.** Quantification of plant height for silenced plants and the vector control using box-and-whisker plots with individual data points (blue, gray and orange data points represent three biological replicates of nine plants each); the boxes show the 25th and 75th percentiles, the median is indicated by a horizontal line and the minimum and maximum values by the ends of the whiskers. *P<0.0001.

**Supplemental Table S1: Transformants used in this study**

| Strain | Used In: | Effector:FP | Description (background strain; plasmid) |
|---|---|---|---|
| KV168 | Fig.1B; Sup. Fig. S2 | Pwl2:mCherry:NLS & Bas4:eGFP | Guy11; pBV591 (Hyg$^R$) |
| KV170 | Fig. 2B; Sup. Fig. S1; Sup. Fig. S6 | Bas1:mRFP | Guy11; pBV440 (Gen$^R$) |
| KV174 | Sup. Fig. S1 | Pwl1:eGFP | Guy11; pBV249 (Hyg$^R$) |
| KV176 | Fig. 3D,E; Fig. 5C; Sup. Movie S2 | Pwl2:eGFP | Guy11; pBV252 (Hyg$^R$) |
| KV182 | Sup. Fig. S1 | Bas1:eYFP | Guy11; pBV231 (Hyg$^R$) |
| KV209 | Fig. 3A-C; Fig. 4A-D; Fig. 5A,B; Fig. 6A-D; Fig. 7D-F; Fig. 9A-E; Sup. Movie S3 | Pwl2:mRFP | Guy11; pBV1192 (Gen$^R$) |
| KV211 | Fig. 2A-F; Sup. Movie S1 | Pwl2:mRFP and Bas1:eYFP | Guy11, pBV231 (Hyg$^R$); pBV1192 (Gen$^R$) |
| KV217 | Fig. 1A,C; Fig. 7C; Fig. 8A-C | Pwl2:mRFP & Bas4:eGFP | Guy11, pBV436 (Hyg$^R$) |
| KV220 | Fig. 4B,C | Bas83:mRFP | Guy11, pBV1194 (Gen$^R$) |
| KV222 | Fig. 4A; Sup. Fig. S4A | Bas83:mRFP & Pwl2:eGFP | Guy11, pBV252 (Hyg$^R$); pBV1194 (Gen$^R$) |
| KV224 | Fig. 1D-F; Sup. Fig. 2; Sup. Fig. S6 | Bas170:mRFP | Guy11, pBV1196 (Gen$^R$) |
| KV244 | Sup. Fig. 1 | Pwl1:mRFP | Guy11, pBV1211 (Gen$^R$) |
| KV246 | Sup. Fig. S4B | Bas83:mRFP & Pwl1:eGFP | Guy11; pBV249 (Hyg$^R$); pBV1194 (Gen$^R$) |

**Supplemental Table 2:** Key plasmids used in this study

| Clone | Description |
|---|---|
| pBV231 | *BAS1* (MGG_04795.6) promoter and entire coding sequence with a C-terminal translational fusion of the eYFP reporter gene. A 1.3-kb PCR product containing the *BAS1* gene was amplified by PCR, digested with *Eco*RI and *Bam*HI, and subsequently cloned in *Eco*RI-*Bam*HI sites of pBV181 containing eYFP in pBHt[2] (AddGene Plasmid #104175). (Hygromycin$^R$, Kanamycin$^R$) Published in Mosquera et al. (2009). *Used in transformants KV182 and KV211.* |
| pBV249/ pSK1898 | *PWL1* (GenBank:U36923.1) promoter and entire coding sequence with a C-terminal translational fusion of the eGFP reporter gene. A 1,171-bp *Kpn*I-*Bam*HI fragment of pSK1876 ($P_{PWL1}$:PWL1CDS) and the 986-bp *Bam*HI-*Xba*I fragment of pSK1873 (eGFP:Ter) cloned between the *Kpn*I and *Xba*I sites of pBHt2. Plasmid published in Khang et al. (2010). *Used in KV174 and KV246.* |
| pBV252/ pSK1901 | *PWL2* (MGG_04301.6) promoter and entire coding sequence with a C-terminal translational fusion of the eGFP reporter gene. Plasmid published in Khang et al. (2010). *Used in transformants KV176 and KV222.* |
| pBV367 | mRFP binary expression vector derived from pBGt (Seogchan Kang, Pennsylvania State University), consisting of three modules: P27 promoter (*Eco*RI-*Bam*HI fragment), mRFP (*Bam*HI-*Sph*I fragment), and *N. crassa* β-tubulin terminator (*Sph*I-*Hind*III fragment) cloned in *Eco*RI-*Hind*III sites of pBGt. (Geneticin$^R$, Kanamycin$^R$) Plasmid published in Giraldo et al. (2013). |
| pBV436 | Plasmid expressing $P_{PWL2}$:PWL2CDS:mRFP:Ter and $P_{BAS4}$:BAS4CDS:eGFP:Ter cloned in the *Eco*RI and *HindIII* sites of pBHt2 (AddGene #104175). Published in Khang et al (2010). *Used in KV217.* |
| pBV440 | *BAS1* (MGG_04795.6) promoter and entire coding sequence with a C-terminal translational fusion of the mRFP reporter gene. Plasmid published in Mosquera et al. (2009). *Used in transformant KV170*. |
| pBV591 | Used to express Pwl2:mCherry:NLS (with an added nuclear localization signal) together with Bas4:eGFP. ProPWL2:PWL2CDS:mCherry:NLS:Tnos and ProBAS4:BAS4CDS:eGFP:Ter cloned in *Eco*RI and *Hind*III sites of pBHt2. (Hygromycin$^R$). Published in Khang et al. (2010). *Used in KV168.* |
| pBV1192 | *PWL2* (MGG_04301.6) promoter and entire coding sequence with a C-terminal translational fusion of the mRFP reporter gene. A 1.7-kb *Pwl2* gene fragment cloned into *EcoR*I-*BamH*I restriction sites of pBV367 (Geneticin$^R$, Kanamycin$^R$). *Used in transformants KV209 and KV211.* |
| pBV1194 | *BAS83* (MGG_08506.6) promoter and entire coding sequence with a C-terminal translational fusion of the mRFP reporter gene. A 1.6-kb *BAS83* gene fragment cloned into *EcoR*I-*BamH*I restriction sites of pBV367. (Geneticin$^R$, Kanamycin$^R$). *Used in transformants KV220, KV222 and KV246.* |

| | |
|---|---|
| pBV1196 | *BAS170* (MGG_07348.6) promoter and entire coding sequence with a C-terminal translational fusion of the mRFP reporter gene. A 2.0-kb *BAS170* gene fragment cloned into the *EcoR*I restriction site of pBV1214. (Geneticin$^R$, Kanamycin$^R$). *Used in transformant KV224.* |
| pBV1198 | Clathrin-mediated endocytosis (CME) fluorescent marker for expression in rice. A 4.71-kb fragment of *Clathrin Light Chain*-1 (*CLC1*) (promoter and entire ORF of rice *CLC1* gene; LOC4337419) integrated into *Kpn*I and *Xho*I restriction sites of pSH1.6_EGFP (AddGene #42323). A 5.7-kb CLC1:eGFP construct was PCR amplified and integrated into pENTR (pENTR™/D-TOPO™ Cloning Kit, ThermoFisher Scientific) and transferred to *Agrobacterium* vector pIPKb001 (Himmelbach et al., 2007) using the Gateway LR Clonase II system. *Used to express CLC1:eGFP in rice cv. YT16.* |
| pBV1200 | Clathrin-independent endocytosis (CIE) fluorescent marker for expression in rice. A 4.4-kb fragment of *Flotilin 1* (promoter and entire ORF of rice *Flot1* gene; LOC4348926) integrated into *Hind*III and *Eco*47III restriction sites of pSH1.6_EGFP (AddGene #42323). A 5.4-kb Flot1:eGFP construct was PCR amplified and integrated into pENTR (pENTR™/D-TOPO™ Cloning Kit, ThermoFisher Scientific) and transferred to *Agrobacterium* vector pIPKb001 (Himmelbach et al., 2007) using the Gateway LR Clonase II system. *Used to express Flot1:eGFP in rice cv. YT16.* |
| pBV1202 | VIGS vector for silencing the *Flot1* gene in rice cv. IR64. A 0.3-kb fragment for targeted silencing of the rice gene for CIE component Flotillin-1 (*Flot1* gene; LOC4348926) was cloned in *Avr*II-*Nco*I restriction sites of the pC13/F3-13m VIGS vector (Kanamycin$^R$) as described in Ding et al. (2006 and 2007). |
| pBV1204 | VIGS vector for silencing the *AP-2α* gene in rice. A 0.3-kb fragment for targeted silencing of rice gene for CME component *Adaptor Protein Complex-2α* (*AP-2α* gene; LOC4331370) was cloned into *Avr*II-*Nco*I restriction sites of the pC13/F3-13m VIGS vector (Kanamycin$^R$) (Ding et al. 2006, 2007). |
| pBV1206 | VIGS vector for silencing the *CHC1* gene in rice cv. IR64. A 0.3-kb fragment for targeted silencing of the rice gene for CME component *Clathrin Heavy Chain-1* (*CHC1* gene; LOC4349546) was cloned in *Avr*II-*Nco*I restriction sites of the pC13/F3-13m VIGS vector (Kanamycin$^R$) as described in Ding et al. (2006 and 2007). |
| pBV1211 | *PWL1* (GenBank:U36923.1) promoter and entire coding sequence with a C-terminal translational fusion of the *mRFP* reporter gene. A 1.5-kb *PWL1* gene fragment cloned into the *Bam*HI restriction site of pBV367. (Geneticin$^R$, Kanamycin$^R$). *Used in transformant KV244.* |
| pBV1214 | mRFP binary expression vector derived from pBGt (Seogchan Kang, Pennsylvania State University), consisting of four modules: P27 promoter (*Eco*RI-*Bam*HI fragment), multiple cloning site from pSH1.6_EGFP containing an extra *Eco*RI site cloned into the *Bam*HI restriction site, mRFP (*Bam*HI-*Sph*I fragment), and *N. crassa* β-tubulin terminator (*Sph*I-*Hind*III fragment) cloned in *Eco*RI-*Hind*III sites of pBGt. (Geneticin$^R$, Kanamycin$^R$). *Used in KV224.* |

**Supplemental Table 3:** Oligonucleotides used in this study

| Primer name | Sequence 5' -3' | Description |
|---|---|---|
| **Primers used to fluorescently label effectors in *M. oryzae*** | | |
| MoPLW2-EcoRI-F1* | AAAGAATTCTTACCCGTGGCAAGGATAAC | 1.6-kb *Pwl2* promoter plus entire coding sequence |
| MoPLW2-BamHI-R1 | TTTGGATCCCATAATATTGCAGCCCTCTTCTCG | |
| MoPLW1-BanHI-F1 | AATGGATCCGCCTGTCACTAGGGCTTCTG | 1.5-kb *Pwl1* promoter plus entire coding sequence |
| MoPLW1-BamHI-R1 | ATTGGATCCCATAATTGGCAGCCCTGATCTC | |
| mRFPORF1-R1 | CTCCTCGCCCTTGCTCACCAT | 1.5-kb test PCR to check the orientation of *PWL1* in pBV367 |
| MoBas83EcoRI-F1 | ATAGAATTCAGAGTGTGCGCTAGCTCGAC | 2.0-kb *Bas83* promoter plus entire coding sequence |
| MoBas83BamHI-R1 | TATGGATCCACCCGTCCGCTGCGAAGTC | |
| MoBas170EcoRI-F1 | AAAGAATTCTCGTTGCAGCGGCAGTCAAG | 2.0-kb *Bas170* promoter plus entire coding sequence |
| MoBas170EcoRI-R1 | AAAAGAATTCGCCGTCTCCACAGCCCTGGAT | |
| MoBas170ORF-F1 | GTGTCGTGGTTAGGACAATG | 0.772-kb test PCR to check the orientation of *Bas170* in pBV367; used with the primer mRFPORF1-R1 |
| **Primers used to gene replacement in *M. oryzae*** | | |
| M13F | CGCCAGGGTTTTCCCAGTCACGAC | 1.113-kb of HY split cassette (Catlett et al., 2003) |
| HYsplit | GGATGCCTCCGCTCGAAGTA | |
| M13R | AGCGGATAACAATTTCACACAGGA | 0.74-kb of YG split cassette (Catlett et al., 2003) |
| YGsplit | CGTTGCAAGACCTGCCTGAA | |
| Bas83:BASKOtest-F1 | ATGCGATTCTCGACCGTTTTC | 0.603-kb of the *Bas83* ORF (test PCR to verify the presence of *Bas83* gene in the transformants |
| Bas83:BASKOtest-R1 | ACCCGTCCGCTGCGAAGTC | |
| Bas83LF-F | AGAGTGTGCGCTAGCTCGAC | 1.0-kb 5'-flanking region of MGG_08506.6; used with the split marker method |
| Bas83LFHY-R | *GTCGTGACTGGGAAAACCCTGGCG*GCAAGGCGGCCAGGACAAC | |
| Bas83RFYG-F | *TCCTGTGTGAAATTGTTATCCGCT*ACGCGTCGTTGTAGTGACTTG | |

| | | |
|---|---|---|
| Bas83RF-R | TAACAGCAGTCTGCCCAACAC | 1.0-kb 3'-flanking region of MGG_08506.6; used with the split marker method |
| Bas83LFHYG-R | *CCTCCACTAGCTCCAGCCAAGCC*GCAAGGCGGCCAGGACAAC | 1.0-kb 5'-flanking region of MGG_08506.6; used with the primer Bas83LF-F |
| Bas83LFHYG-F | *TAGAGTAGATGCCGACCGCGGGTT*ACGCGTCGTTGTAGTGACTTG | 1.0-kb 5'-flanking region of MGG_08506.6; used with the primer Bas83RF-R |
| HYG-1F | GGCTTGGCTGGAGCTAGTGGAGG | 1.4-kb hygromycin gene |
| HYG-2R | AACCCGCGGTCGGCATCTACTCTA | |
| **Primers used to silence endocytic machinery in rice** | | |
| OsFlot1AvrII-F1 | AAACCTAGGGTGTATATTGCTGTGTAACTG | 0.3-kb *Flot1* RNAi sequence target |
| OsFlot1NcoI-R1 | TTTCCATGGTGCAGTTATATACTCATGTAAATC | |
| OsClat1HeavyAvrII-F1 | AAACCTAGGAGGAGGCATCTTTCAAGTTGTAG | 0.3-kb *CHC1* RNAi sequence target |
| OsClat1HeavyNcoI-R1 | TTTCCATGGGACATGAGACCGTAAACAAGTAAC | |
| OsAP-2AvrII-F1 | AAACCTAGGGCGGAGCTTTCTTTCTAG | 0.3-kb *AP-2α* RNAi sequence target |
| OsAP-2NcoI-R1; | TTTCCATGGGAATTCACCCATCAATTTAATAAG | |
| OsFlotillin1-ORF-F1 | ATCAACGCCGACGCCATCAG | qRT-PCR primers for rice *Flot1* |
| OsFlotillin1-ORF-R1 | GGCGGCAGCATCTTGTACAC | |
| OsAP2a-ORF-F1 | GAACAGGCCGCCAATTCAG | qRT-PCR primers for rice *AP-2α* |
| OsAP2a-ORF-R2 | TGATCCAGCCCTTGTGATGTAG | |
| OsCHC1-ORF-F1 | ATGCCCAATTGCTTCCTCTC | qRT-PCR primers for rice *CHC1* |
| OsCHC1-ORF-R2 | ATCCCATATGCTGGCATAGG | |
| EF F2 | CCGCCAAGAAGAAATGAGCA | qRT-PCR primers for rice *EF-1α* gene (Ding et al., 2006) |
| EF R2 | TCCATGCAACGAGTGCCAT | |
| **Primers used to fluorescently label endocytosis markers in rice** | | |
| KpnI_OsCLC1Prom1-F1 | TTGGTACCCTGGAGAGCCCAGGCATTAC | 4.71-kb Promoter and entire ORF of rice *CLC1* gene |

| | | |
|---|---|---|
| XhoI_OsCLC1-R1 | CGCTCGAGGCTCCGATGCTGCAGGCTGCTC | |
| CACC_OsCLC1-F1 | CACCCTGGAGAGCCCAGGCATTAC | 5.657-kb CLC1:eGFP |
| eGFPT2-R2 | TGAAGGCGTACTAGGTTGCAGTC | |
| HindIII-OsFlot1Prom-F1 | AAAAGCTTCAAAGCCGCTAACGGTTTGG | 4.424-kb Promoter and entire ORF of rice *Flot1* gene |
| Eco47III_OsFlot-R1 | AAAAGCGCTGGACTGGTCGACGAGGGGGCGC | |
| CACC_OsFlot1Prom1-F1 | CACCCAAAGCCGCTAACGGTTTGG | 5.367-kb Flo1:eGFP; used with the primer eGFPT2-R2 |

*Restriction sites are underlined

**Legend for Supplemental Movie S1**

**Optical sections showing a cytoplasmic connection between a side-BIC and peripheral rice cytoplasm as well as effector vesicles in different cell layers.** This movie documents optical sections down through a leaf sheath cell invaded by strain KV211 (secreting Pwl2:mRFP and Bas1:eYFP) at 24 hpi. Effector vesicles, the cytoplasmic connection, and the host cytoplasm are labeled with cytoplasmic effectors Pwl2:mRFP and Bas1:eYFP. Pwl2:mRFP fluorescence (magenta) appears to predominate at this point, which is consistent with the strong Pwl2:mRFP fluorescence at 21 hpi in Figure 1A. One optical section showing the cytoplasmic connection is reproduced below. The file shows merged bright field, mRFP and eGFP.

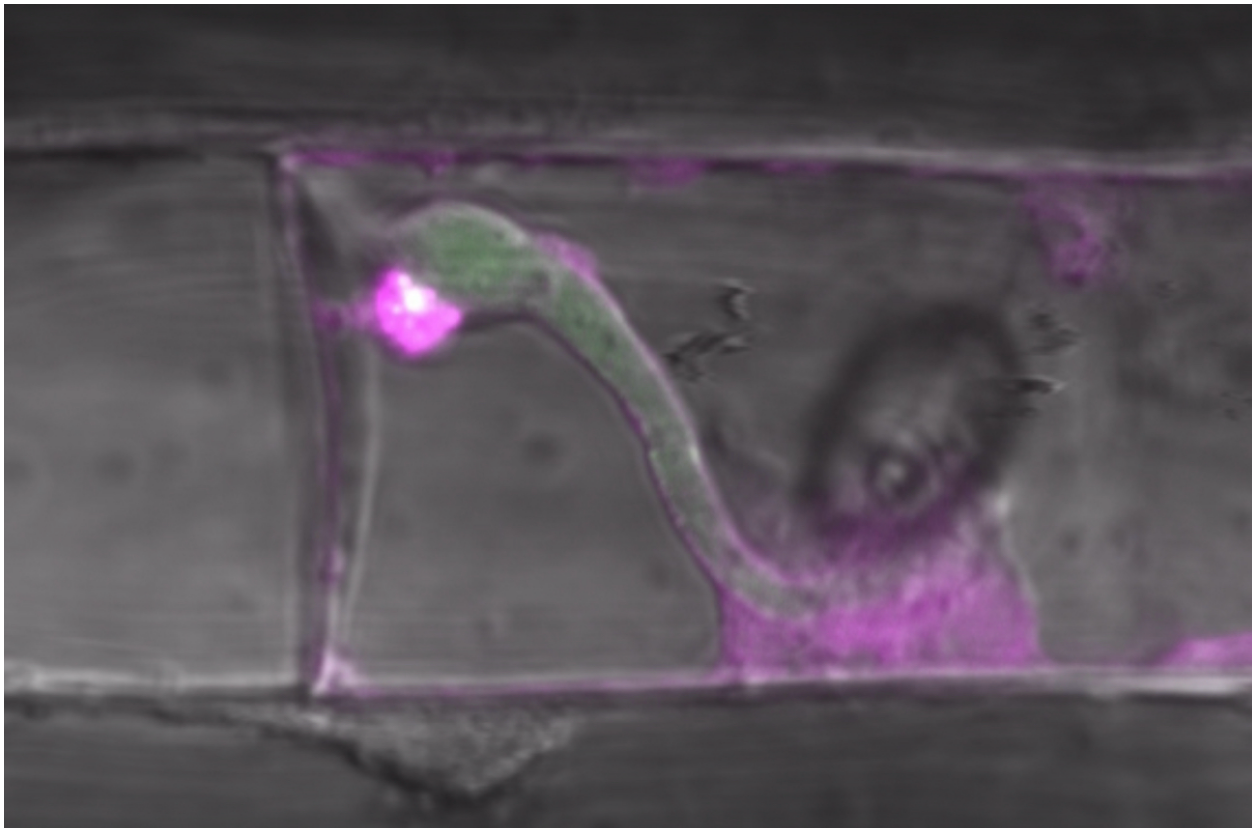

**Legend for Supplemental Movie S2**

**Optical sections showing effector vesicles in the host cytoplasm and at a distance from a BIC, in rice tissue stained by endocytosis tracer dye FM4-64.** Vesicles are visualized by Pwl2:eGFP, FM4-64 or both. The lower section from the movie shows FM4-64 tightly encasing the invasive hypha (strain KV176), but excluded from fungal septa and vacuolar membranes. This indicates the EIHM is intact around the IH and protecting fungal membranes from insertion of the FM4-64 dye. The image below is a single optical section from the movie. The file shows merged mRFP (magenta) and eGFP (green).

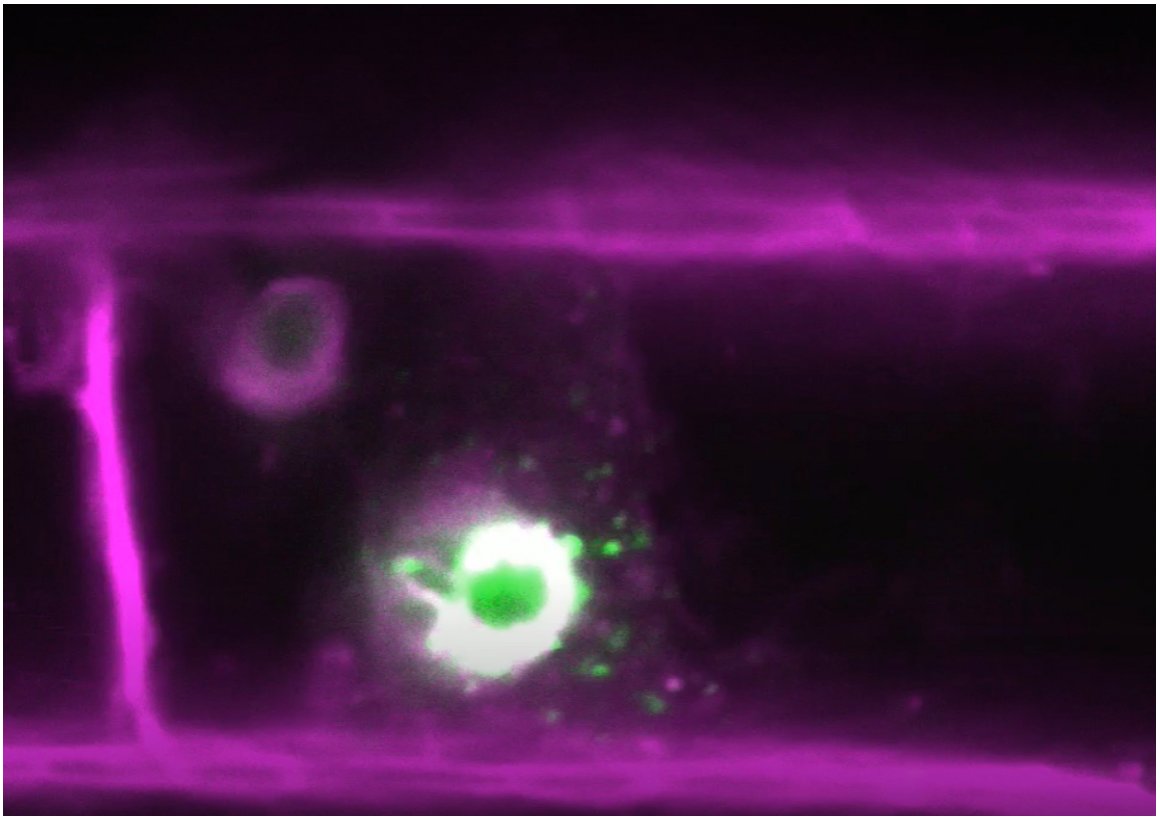

**Legend for Supplemental Movie S3**

**Time-lapse images showing co-localization of clathrin light chain-1 (CLC-1:eGFP) labeled with eGFP and cytoplasmic effector Pwl2 (Pwl2:mRFP) labeled with mRFP in BIC vesicles.** Time lapse images of a KV209 invasive hypha secreting Pwl2:mRFP and growing in CLC1:eGFP-labeled rice immediately after differentiation of the primary hypha into a bulbous IH cell. In the movie image below, compare white in the side-BIC vesicles due to co-occurrence of red cytoplasmic effector fluorescence and green CLC-1:eGFP fluorescence with the green CLC1:eGFP foci around cell periphery and under the appressorium. The file contains merged bright field, mRFP (magenta) and eGFP (co-localization appears white).

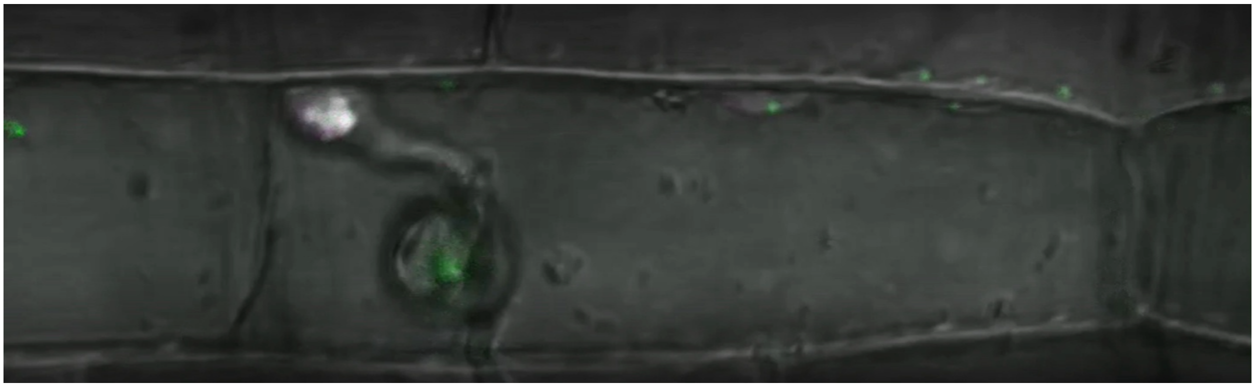
